# Divergent age-related methylation patterns in long and short-lived mammals

**DOI:** 10.1101/2022.01.16.476530

**Authors:** Amin Haghani, Nan Wang, Ake T. Lu, Khyobeni Mozhui, Mammalian Methylation Consortium, Ken Raj, X. William Yang, Steve Horvath

## Abstract

Age-related changes to cytosine methylation have been extensively characterized across the mammalian family. Some cytosines that are conserved across mammals exhibit age-related methylation changes that are so consistent that they were used to successfully develop cross-species age predictors. In a similar vein, methylation levels of some conserved cytosines correlate highly with species lifespan, leading to the development of highly accurate lifespan predictors. Surprisingly, little to no commonality is found between these two sets of cytosines even though the relationship between aging and lifespan is by most measures linked. We ventured to address this conundrum by first identifying age-related cytosines whose methylation levels change in opposite directions between short and long-lived species. We hypothesized that age-related CpGs that are also associated with species lifespan would tap into biological processes that simultaneously impact aging and lifespan. To this end, we analyzed age-related cytosine methylation patterns in 82 mammalian species. For each CpG, we correlated the intra-species age correlation with maximum lifespan across mammalian species. We refer to this correlation of correlations as *Lifespan Uber Correlation (LUC)*. This approach is unique in incorporating age and species lifespan in a single analysis. We identified 629 CpGs with opposing methylation aging patterns in long and short-lived species. Many of these are found to be near *BCL11B*, *NPTN,* and *HOXC4* loci. Methylation and transcription analyses of *Bcl11b* knockout mice indicate that this gene partially regulates the methylation state of LUC CpGs. We developed DNAm age estimators (epigenetic clocks) based on LUC CpGs. These LUC clocks exhibited the expected behavior in benchmark aging interventions such as caloric restriction, growth hormone receptor knockout and high-fat diet. Furthermore, we found that *Bcl11b* heterozygous knockout mice exhibited an increased epigenetic age in the striatum. Overall, we present a bioinformatics approach that identified CpGs and their associated genes implicated both in aging and lifespan. These cytosines lend themselves to developing epigenetic clocks that are sensitive to perturbations that impact both age and lifespan.

## INTRODUCTION

Aging is partly a result of the gradual accumulation of molecular damage that increasingly compromises tissue and organ function. The cause of such stochastic damage and its persistence is a complex mix of various stressors ^1^. In contrast, maximum lifespan appears to be an inborn species trait that reflects, in part, the innate capacity of an organism to protect against damage from the environment. As such, maximum lifespan is deemed to be a stable trait in species. In contrast, average lifespan results from genetic variations, environmental conditions or stochastic differences encountered by individuals within a species. For example, the maximum human lifespan is set at 122 years by Jeanne Calment, the oldest documented human to date ^2^, while the average human lifespan is 72.6 years. Some studies suggest that the increase in human lifespan started to plateau after 1980 ^3^. Although debatable, human lifespan appears to be reaching its biological maximum. Several evolutionary theories, including antagonistic pleiotropy and the disposable soma theory, were proposed to explain how different maximum lifespans evolved in animals. It is proposed that low extrinsic mortality (predation), an abundance of food, and low reproductive cost are features that offer long-lived species the opporunity to evolve better or more efficient mechanisms to protect against age-mediated accumulation of damage. As such, the rate of biological function decline (i.e., aging) would be expected to correlate inversely with maximum species lifespan. Although aging and maximum lifespan are intimately intertwined, they nevertheless appear in some biological investigations to be distinct processes ^4^. One of the most direct ways to investigate this conundrum is through comparative studies between mammalian species. The mammalian family is constituted of species with over a hundred fold range in maximum lifespan and 50 million fold differences in body size and weight. The latter is an important consideration, given that there is a very high correlation between species weight and lifespan.

To address the aging-lifespan link, we leveraged the DNA methylation database that was generated by the Mammalian Methylation Consortium. This database houses cytosine methylation profiles from hundreds of mammalian species, and has already been employed to characterize the relationship between CpG methylation and chronological age^5–25^, and to identify cytosines related to life-history traits such as maximum lifespan and gestation time^15, 26^. These analyses did indeed uncover the relationship between mean methylation levels of some cytosines conserved across species to maximum species lifespan^15^, but surprisingly these CpGs tended to be different from those that correlate strongly with chronological age^15, 16^. While this lack of overlap can be explained on statistical grounds (different regression models) and biological differences (inborn traits vs individual characteristics), it runs counter to the intuition that maximum lifespan reflects intrinsic mortality risk associated with biological aging. Here we pursue a different approach, one that is based on the hypothesis that age-related cytosines that change their methylation states in opposing directions in long and short-lived species are the biological linchpins that link aging with lifespan. We reason that the methylation states of these cytosines are more likely to impose biologically relevant effects that impact both aging as well as lifespan.

Our recent study of bats revealed that age-related CpGs with opposing methylation patterns in long and short-lived bat species are proximal to genes that are related to the immune system and cancer^27^. Here, we expanded this single species study along several lines. First, we profiled 82 mammalian species, including exceptionally long-lived ones such as bowhead whales and naked mole rats, as well as many short-lived species such as mice. Second, we analyzed several tissue types (e.g., blood, skin, muscle, liver). Third, we developed a statistical measure (lifespan uber correlation) to quantify differences in aging patterns between short and long-lived species. Fourth, we studied the impact of adult weight on these aging patterns. Fifth, we used transcriptomic and mouse knockout studies to characterize some of the most significant CpGs. Finally, we used these CpGs to build age estimators (epigenetic clocks) and demonstrated that these clocks are able to capture the effects of interventions that are known to alter age we well as lifespan.

### Lifespan Uber Correlation

We analyzed previously-described data from the Mammalian Methylation Consortium ^15, 16, 26^. We used n=8,500 tissue samples (from blood, skin, liver, cerebral cortex, and muscle) collected at different ages (neonatal to old life-stages) of 82 mammalian species with a diverse range of reported lifespan (3.8 years in rats, to 211 years in bowhead whales). These data were generated using the mammalian methylation array platform that measures methylation levels at highly conserved CpGs across mammalian species ^28^. We restricted our analyses to 14,705 CpGs that are conserved in eutherian species (Methods) ^26^.

We define “*Lifespan Uber Correlation*” (LUC) as the correlation between age-related CpGs and maximum species lifespan. This is derived by correlating the correlation between CpG and chronological age, with maximum lifespan (**Fig. 1b**). Maximum lifespan was first transformed using a base e log transformation. For each CpG and tissue type, the LUC was implemented in two steps. First, the Pearson correlation was determined between the CpG and chronological age for all tissue samples from a given species. This results in a value, F(Species)=cor(CpG,Age), that is specific to the tissue type of the species in question. Second, F(Species) is correlated with log-transformed maximum lifespan across all available species.

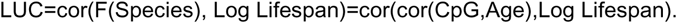

**Fig. 1.**
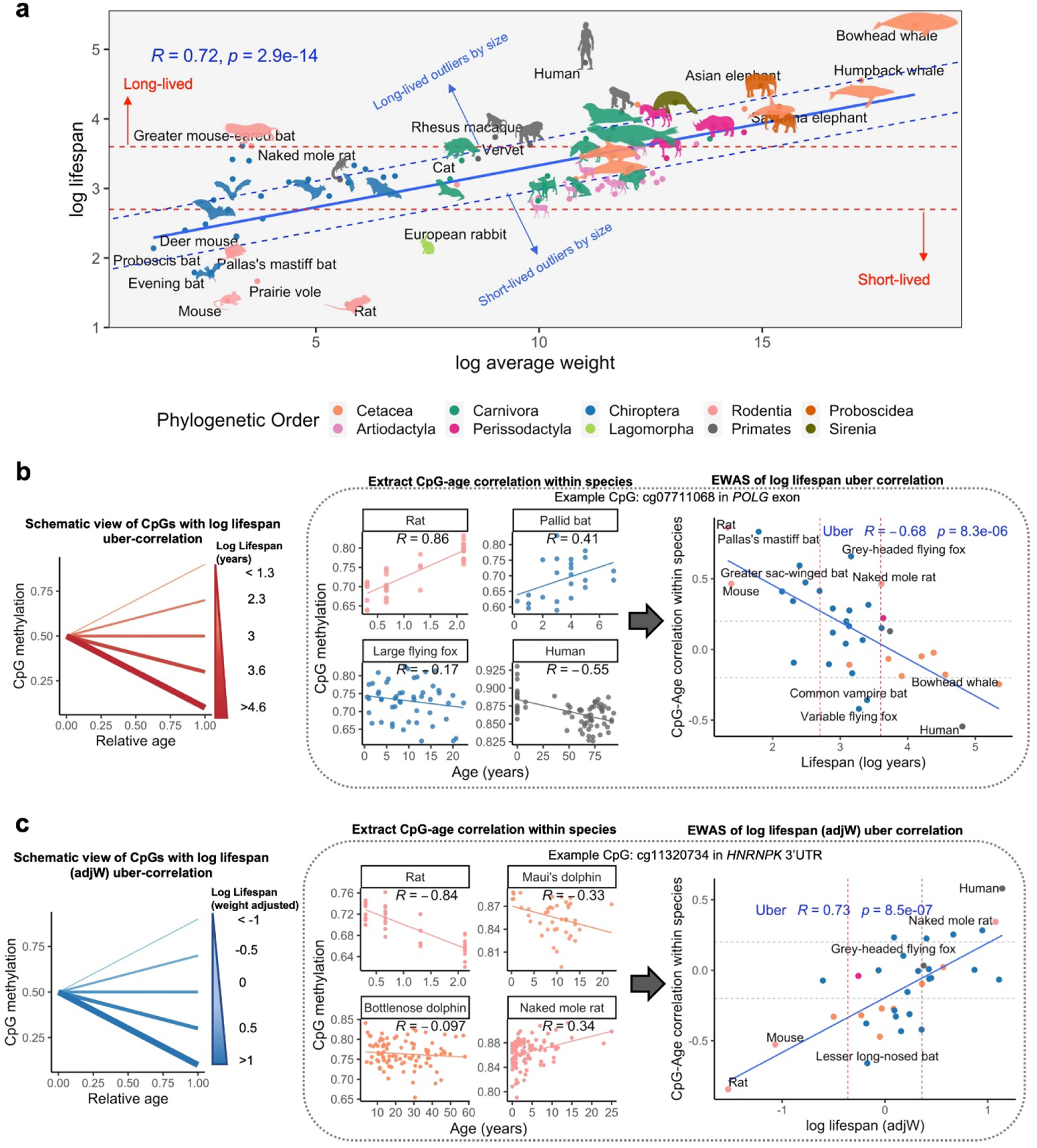
Schematic view of lifespan uber correlation analysis in mammalian data. **a**, Scatter plot of maximum lifespan and average adult weight of the mammalian species in this study. **b**, Schematic chart of the CpGs with significant LUC in mammals. The LUC analysis first correlates intraspecies correlations between a given CpG and chronological age, cor(CpG,Age). Next it forms the inter-species correlation with log-lifespan: LUC=cor(Log Lifespan,cor(CpG,Age)). Only the species with a sample size of >10 (range 11 to 1,111) and at least 20% coverage of lifespan were included in this analysis. The number of species per tissue: Blood, 30 species, Skin, 34 species. **c,** Schematic chart of the CpGs with the uber correlation of log lifespan adjusted for body size (lifespanAdjW). This analaysis includes the inter-species correlation of within species CpG-Age correlations with the residuals of log-lifespan regressed on log-weight of the animals. LUC.adjW=cor(residuals,cor(CpG,Age)). Only the species with a sample size of >10 (range 11 to 1,111) and a 20% coverage of lifespan were included in this analysis. The number of species per tissue: blood, 30 species, skin, 34 species.

Note that the final expression involves a correlation coefficient on top of another. The German word “Uber- (which means over/above) indicates that one correlation is above the other. CpGs with a negative LUC value exhibit age-related loss of methylation in long-lived species but a gain of methylation in short-lived ones (**Fig. 1b**). Conversely, CpGs with a positive LUC value exhibit age-related gain in long-lived species and loss of methylation in short-lived ones.

It is well established that across the mammalian family, lifespan is significantly correlated with the average adult weight of the species. In our analysis of species characteristics in the AnAge data-base, we also observed a strong correlation (cor=0.72, **Fig. 1a**) ^29^. Therefore statistical analyses of maximum lifespan were executed in two ways: one ignored the effect of adult weight, and the other accounted for this potential confounder. We start by describing the analysis that ignored adult weight.

### LUC analysis without adjustment for adult species weight

In our primary analysis, we applied the LUC analysis to two tissues for which a sufficient number of species (n>30) were available: blood (n=31 species) and skin (n=35 species). In secondary analyses, we applied the LUC method to 3 additional tissue types (cerebral cortex, liver, and muscle, **Extended Data Fig. 1, Extended Data Fig. 2**).

**Fig. 2.**
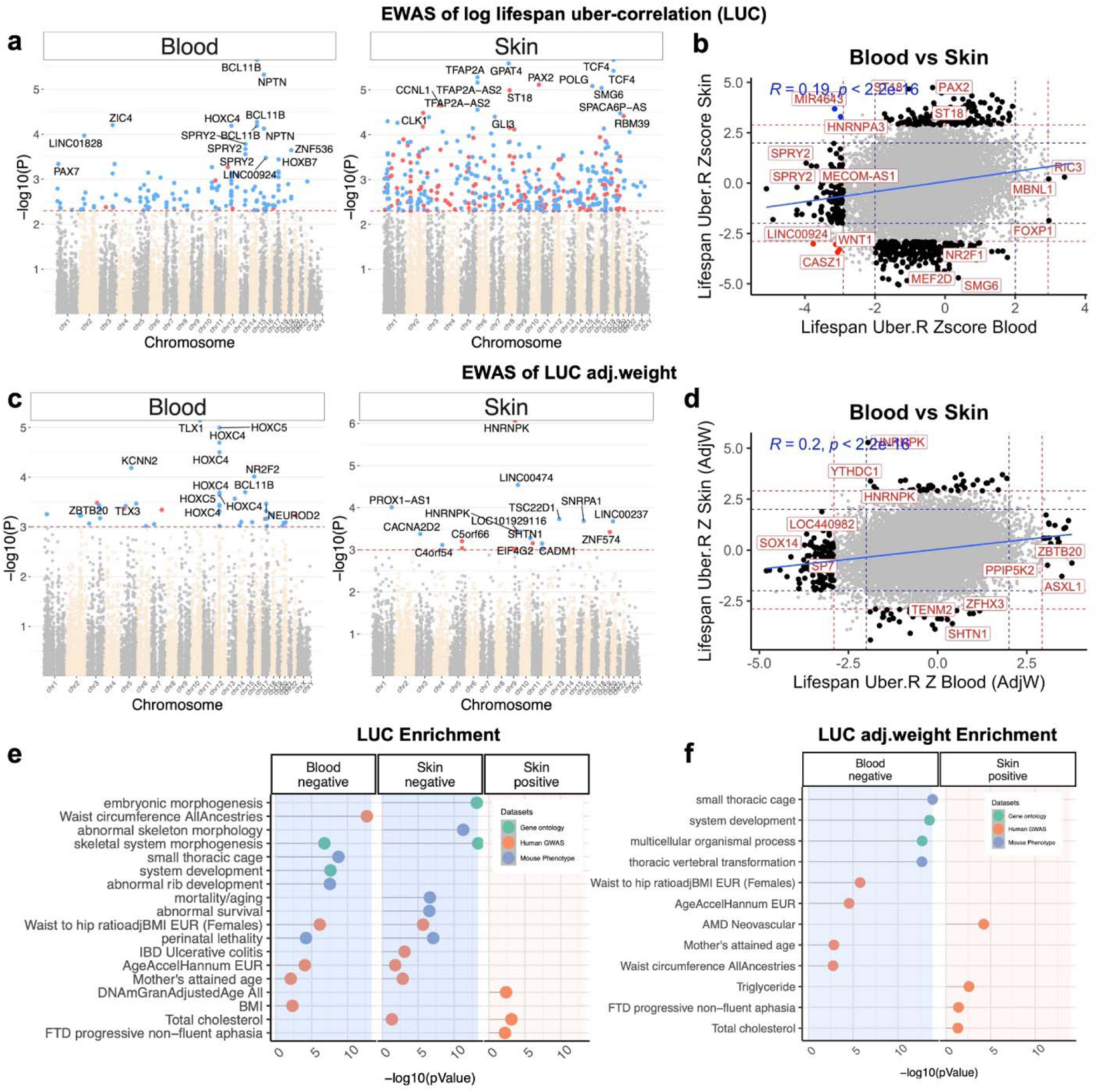
LUC and LUC.adjW EWAS of mammalian skin and blood. **a**, Manhattan plots of the LUC EWAS in 14,705 conserved CpGs in eutherians. The coordinates are based on the alignment to Human hg38 genome. The direction of association is highlighted by red (positive correlation with lifespan) and blue (negative correlation with lifespan) at p<0.005 (red dotted line) significance. **b**, Sector plot of the LUC in skin and blood of mammals. Red-dotted line: p<0.005; blue-dotted line: p>0.05; Red dots: the CpGs that were shared between blood and skin; blue dots: the CpGs with divergent LUC between blood and skin. **c**, Manhattan plots of LUC.adjW analysis in 14,705 conserved CpGs in eutherians. **d**, Sector plot of the LUC.adjW correlations in skin and blood of mammals. **e, f,** Gene set enrichment analysis CpGs with LUC and LUC.adjW correlations.

To assign significance levels to LUC, we applied the Fisher Z transformation to the LUC correlation coefficient resulting in the Uber Z statistic that follows a standard normal distribution under the null hypothesis of no correlation between aging methylation pattern and maximum lifespan. We uncovered 133 CpGs in blood and 433 CpGs in skin that exhibited a significant LUC at a nominal significance level of 0.005, which was selected following a permutation test procedure (**Fig. 2a, Table S1**). The permutation test confirmed that the significance level in the actual data exceeds that in each of the 1000 permutation tests in blood (observed –log10 P = 5.65; maximum permutation log10 P= 4.1 **Extended Data Fig. 3**).

**Fig. 3.**
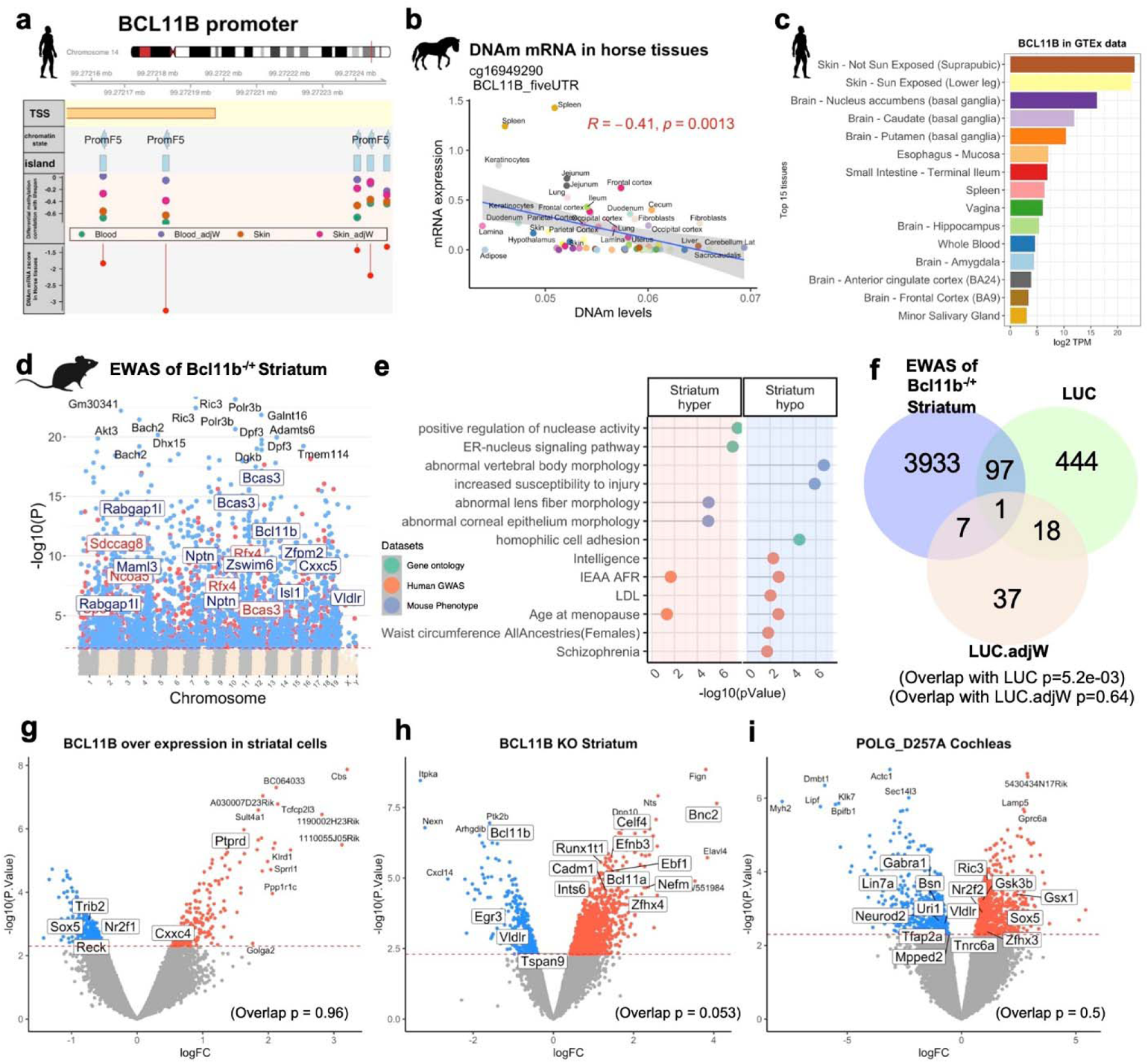
BCL11B partially regulates DNAm and expression of genes with LUC correlation in mammals. **a,** Genome view of BCL11B CpGs with LUC or LUC.adjW correlations. The figure only depicts CpGs that were found to be significant in at least one of the LUC analyses. The genome view plots show the chromosome ideogram, human coordinates (Hg38), Ensemble transcript track, gene-regions annotation of highlighted CpGs, chromatin state of highlighted CpGs, CpG island status of the highlighted CpGs, LUC or LUC.adjW correlation in mammals, and the Z score of DNAm-mRNA association in 29 tissues horses tissues ^31^. **b,** The scatter plot of DNAm-mRNA association in horse tissues were illustrated for a selected CpG in *BCL11B* promoter. **c,** BCL11B has the highest expression in skin and basal ganglia of GTEx data. **d,** EWAS of Bcl11b^−/+^ knock-out striatum. Sample size: 16/group, ages, 0.16-0.5 years. The association was done by a multivariate model with age as co-variate. red line, p<0.005 (FDR<0.03). The results for cerebral cortex and cerebellum is reported in **Extended Data Fig. 10**. The top CpGs overlapping with Uber correlation results are labeled with adjacent genes in the boxes and larger fonts. **e,** Gene set enrichment analysis CpGs related to Bcl11b^−/+^ striatum. **f,** overlap of striatum Bcl11b^−/+^ differentially methylated CpGs with uber correlation hits. Overlap Fisher exact test p = 8.6e-03, Odds ratio = 1.3. **g,h,i,** Transcriptome effects of the Bcl11b overexpression (N=4 per genotype) (**g**), *Bcl11b* heterozygous knockdown (N= 4 *Bcl11b*^+/−^; 3 Control) (**h**), and Polg knockin mutation (N=5 per genotype) (**i**). Datasets: GSE31096, GSE9330, GSE4866. All experimental data have mouse origins. Red line: p<0.005. The genes identified by mammalian LUC and LUC.adjW analyses are highlighted by larger font size and square outline in the figures. The overlap pvalue reports the Fisher exact test of the overlap of differentially expressed genes (p<0.005) with 400 LUC (or LUC.adjW) genes. The background was limited to the genes that are covered in Mammalian Methylation Array.

A high percentage of the significant CpGs had a negative LUC value (95% in blood, 65% in skin, **Extended Data Fig. 1**), indicating that these CpGs lose methylation in long-lived species but gain methylation in short-lived ones. The LUC analysis in blood implicates the following regions: *BCL11B* 5’UTR (Uber Z = −5.1), *NPTN* 2nd exon (Uber Z=−4.9), *ZIC4* 2nd exon (Uber Z=−4.2), and *HOXC4* intron (Uber z=−4.2). The LUC analysis implicated 24 gene regions that contained multiple significant CpGs (e.g., n=4 CpGs for *BCL11B* promoter, n=5 CpGs in the NPTN exon, and n=7 CpGs in the HOXC4 exon), suggesting a robust signal. The LUC analysis in skin implicates *TCF4* 5’UTR (Uber Z=−5.1, n=2 CpGs), *GPAT4* downstream (Uber Z=−5, n=2), *TFAP2A* 2nd intron (Uber Z=*4.8, n=1), and *POLG* 2nd exon (Uber Z=−4.7, n=1; **Fig. 1b**) in skin (**Fig. 2a, Extended Data Fig. 2**).

CpGs that were found in both blood and skin were proximal to *CASZ1* 2nd intron (Blood Uber Z= −3.0, Skin Uber Z= −3.4), *WNT1* downstream (−3.0, −3.3), *LINC00924* 1st intron (−3.7, −3.0), and *POU4F3* 1st exon (−3.1, −3.0), all of which had a negative LUC (**Fig. 2b**). In contrast, two CpGs proximal to *HNRNPA3* upstream (−2.9, 3.2) and *MIR4643* 2nd intron (−3.1, 3.6) had the opposite LUC value between skin and blood, which may be due to biological differences between tissues or may instead reflect false-positive associations.

As a sensitivity test, the LUCs of the top hits were tested in the species that were excluded in the primary analysis due to low sample size. This validation data set had a sample size of 4-10 per species that covered at least 20% of the lifespan. This data set contained 23 species for blood and six species for skin. Interestingly, the LUCs of several top hits such as *BCL11B*, *HOXC4*, *HOXC5*, and *POLG* were corroborated in this validation set (**Extended Data Fig. 4**).

**Fig. 4.**
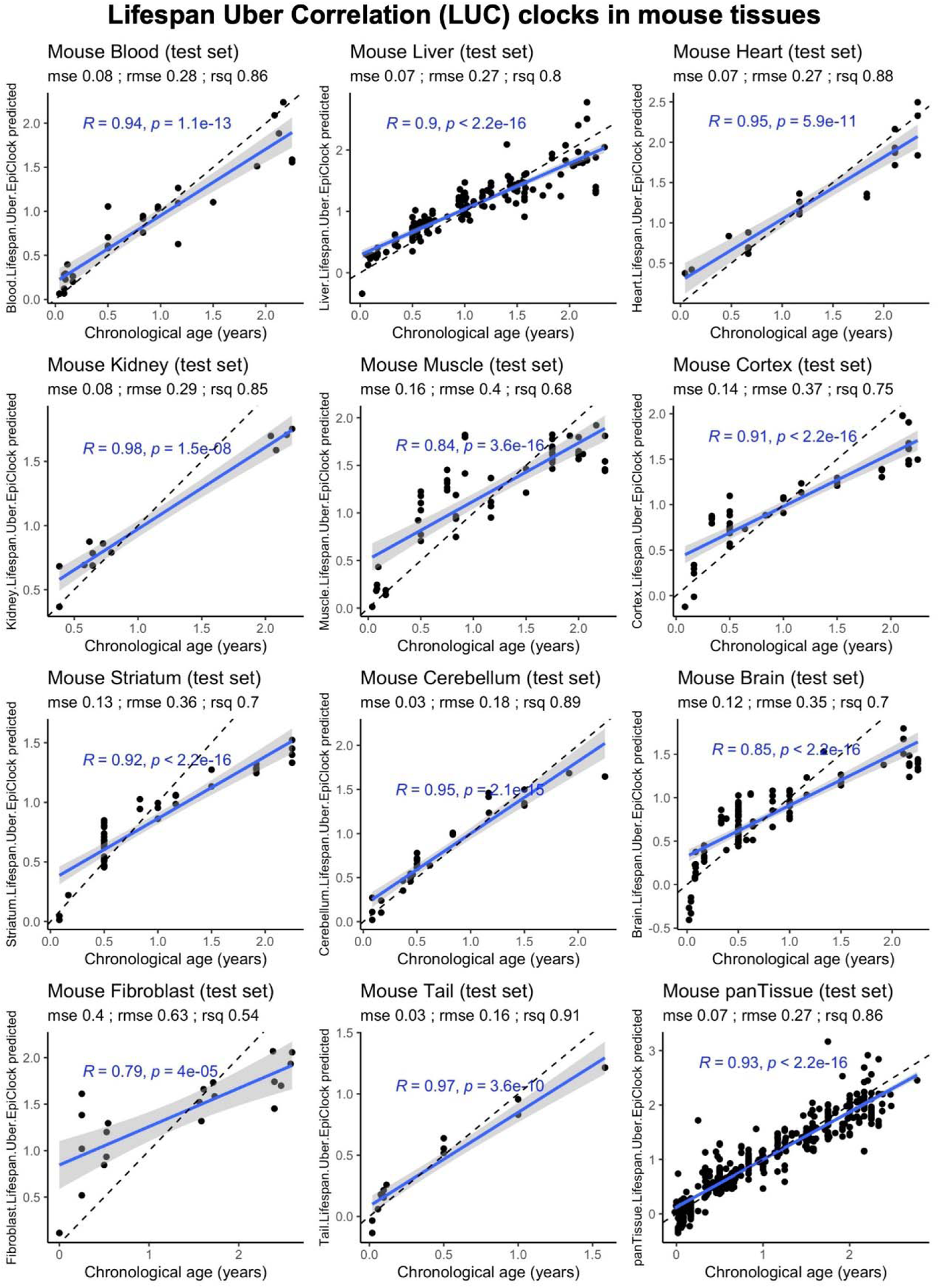
Epigenetic clocks developed from CpGs with significant LUC. The LUC epigenetic clocks are trained in different mouse tissues using 604 CpGs from uber correlation analysis by ridge regression. Total sample size includes: blood, 94; liver, 474; heart, 49; kidney, 45; muscle, 96; cerebral cortex, 185; striatum, 137; panTissue, 720. The samples were split into the train and test sets by 0.7 / 0.3 ratio. The coefficients for the mouse LUC clocks are reported in **Table S6**.

We used the Genomic Regions Enrichment of Annotations Tool (GREAT)^30^ to carry out functional enrichment analyses of the significant CpGs. The CpGs with significant negative LUC values in blood were enriched in loci linked to human waist circumference, as revealed by genome-wide association studies (GWAS) (**Fig. 2e, Extended Data Fig. 5, Table S2**). CpGs with significant negative LUC values in skin, however, appear to be enriched in loci that are implicated in embryonic morphogenesis, mortality/aging, and perinatal lethality. The results for CpGs with positive LUC values were insignificant, which probably reflects the low number of such CpGs.

**Fig. 5.**
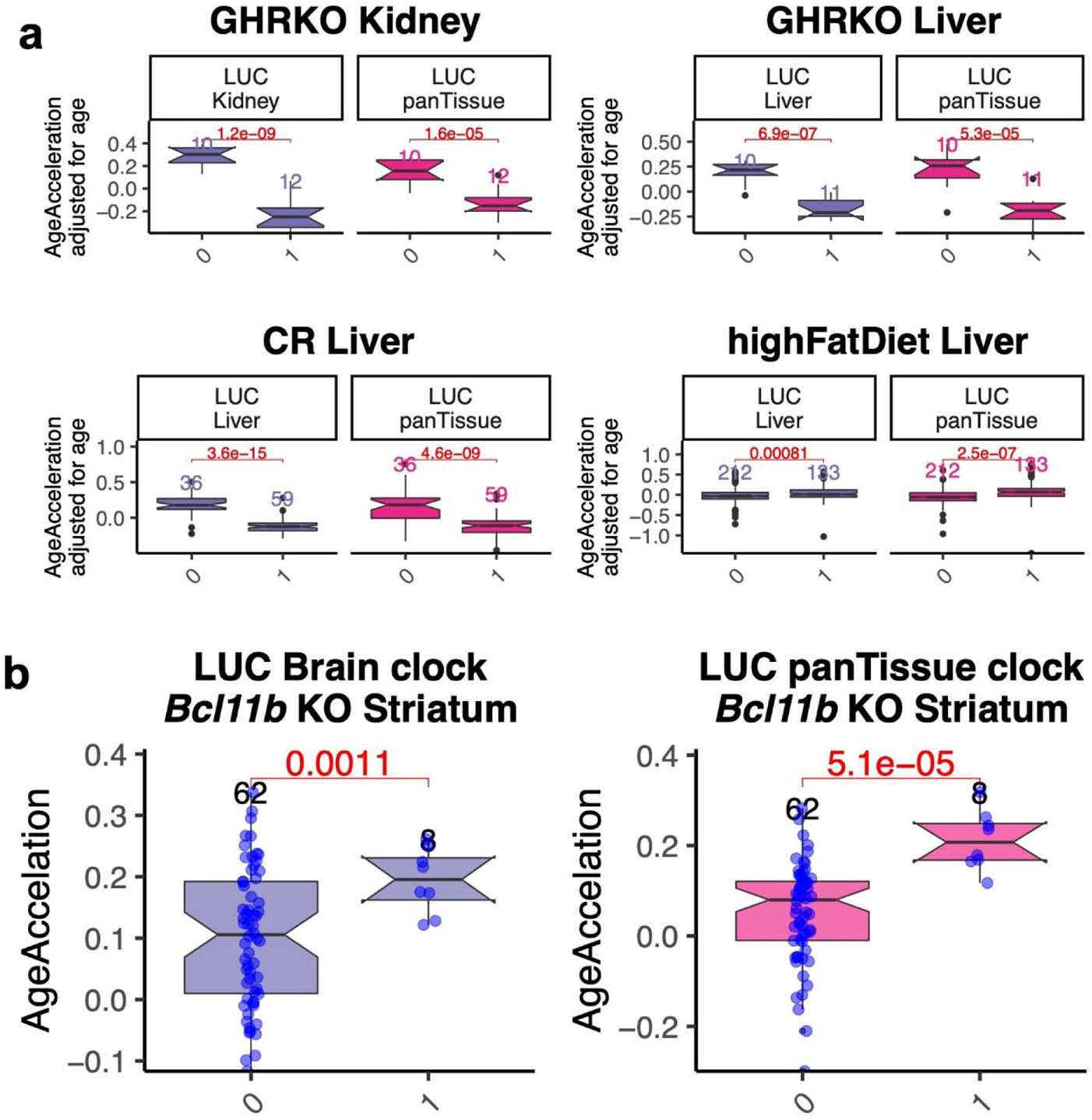
LUC epigenetic clocks are responsive to the lifespan intervention experiments. **a,** The lifespan LUC epigenetic clocks predicts a decrease of epigenetic aging by growth hormone knock out and caloric restriction treatment, but an age acceleration by the high fat diet. **b,** *Bcl11b* knockdown striatum has accelerated aging based on both panTissue and brain LUC clocks that the age matched wildtype mice. “1” indicates the treatment, “0” indicates control (Age matched C57BL6/J mice on normal diet) for each experiment. Reported pvalues in each plot is based on two sided Student T test. The box plots represent lower and upper quartile of the distribution. The notch around the median number of CpGs (horizontal line inside box) depicts the 95% confidence interval. The whiskers extend to the most extreme data point that is no more than 1.5 times the interquartile range from the box. The sample size for each group is reported on top of each box.

### LUC analysis adjusted for adult species weight

To avoid the confounding effect of adult weight, we defined a weight-adjusted measure of maximum lifespan by regressing log lifespan on the log of adult weight and forming residuals. As expected, after adjusting for average adult weight, naked mole rats and humans stand out as long-lived outliers. For example, the naked mole-rat weighs approximately 35 grams but exhibits a maximum lifespan of 37 years ^29^. This number is seven times longer than the lifespan of the comparably-sized prairie voles (40 grams). The weight-adjusted measure of lifespan was used to define another LUC measure for each CpG, denoted as LUC.adjWeight (LUC.adjW) (**Fig. 1c**).

At a nominal value of p<0.001, a total of 44 and 19 CpGs with significant LUC.adjW were identified in blood and skin, respectively. The permutation test of LUC.adjW confirmed that the observed results in blood and skin are statistically significant (**Extended Data Fig. 3, Table S1**). The top regions per tissue are as follows: blood, *TLX1* 2nd exon (Uber Z=−4.8), *HOX5* 1st exon (Z=−4.7), *HOXC4* 5’UTR (−4.5); and skin, *HNRNPK* 3’UTR (Z=5.2), and *LINC00474* upstream (Z=−4.4) (**Fig. 2c, Extended Data Fig. 7**). Similar to LUC analysis (without weight adjustment), multiple significant LUC.adjW CpGs were found on the same gene regions (e.g., *HOXC5*, *HOXC4*, *HNRNPK*). As was observed with LUC, a high percentage of the significant LUC.adjW CpGs exhibited negative values (86% in blood, 58% in skin, **Extended Data Fig. 7**).

The LUC.adjW for blood and skin only exhibited a weak correlation (r=0.2) with each other, and none of the CpGs exhibited significant LUC.adjW values in both tissues at a significance value of 0.001 (**Fig. 2d**). The genes adjacent to significant LUC.adjW CpGs were enriched in developmental processes and implicated in the maternal age of individuals (mother’s attained age). These are similar to the lifespan uber analysis that did not adjust for adult weight (**Fig. 2f, d, Extended Data Fig. 6**). Only a subset of 11 CpGs (25%) in blood (proximal to *BCL11B, SP7*, *NPTN*, *HOXC4*, *HOXC5*, *TLX1*, *LOC440982*, *NR2F2*, *CDKN3*) and 5 CpGs (26%) in skin (proximal to *MSANTD2*, *ZNF574*, *TSC22D1*, *SNRPA1*, *LINC00474*) were shared between LUC correlation analysis of lifespanAdjW and lifespan. It is interesting that methylation changes of CpGs in *BCL11B* were found to be significant in both LUC and its weight-adjusted LUC.adjW counterpart.

### Methylation versus mRNA levels across horse tissues

The identification of cytosines that interface age and lifespan raises the question of whether their methylation level changes lead to gene expression changes. We have access to datasets from horse tissues with which this question can be addressed. We observed that the methylation of several CpGs identified by LUC analysis do indeed correlate significantly with the expression of adjacent mRNA levels across 29 different horse tissues^31^ (**Extended Data Fig. 8**). For example, methylation of CpGs near *BCL11B* promoter exhibited a negative correlation with the expression of this gene in horse tissues^31^ (**Fig. 3b**). Even more specifically, methylation of the most significant CpG in blood, cg16949290, located in the 5’ untranslated region of *BCL11B*, correlates inversely (R= −0.41, p=0.001) with *BCL11B* mRNA levels across all 29 horse tissues.

We used publicly available human and mouse transcriptomic data sets to study aging effects on mRNA levels from select genes implicated by LUC analysis (**Extended Data Fig. 9**).

### Epigenetic studies in Bcl11b heterozygous knockout mice

BAF Chromatin Remodeling Complex Subunit *BCL11B* is a gene located on human chromosome 14 (**Fig. 3a**) and on mouse chromosome 12. Several CpGs in *BCL11B* promoter had a negative LUC and LUC.adjW relationship in both blood and skin of mammals (**Fig. 3a**). Since *BCL11B* is highly expressed in the brain (**Fig. 3c**), we analyzed DNAm levels in several brain regions (striatum, cerebral cortex, and cerebellum) of *Bcl11b*^−/+^ mouse lines (aged between 2 and 6 months). Strikingly, 10 out of 31 CpGs adjacent to the *Bcl11b* gene itself showed a significant methylation change (p<0.005, FDR<0.03) in all considered brain regions (**Extended Data Fig. 10a**). Among those 10 CpGs were the two with highly significant LUC scores in mammalian blood (**Fig. 2a**). At a significance level of p<0.005 (FDR<0.03), the *Bcl11b* knockout caused methylation changes in 4,038, 401, and 221 CpGs in the striatum, cerebral cortex, and cerebellum, respectively (**Fig. 3d; Extended Data Fig. 10a; Table S3**). The large number of differentially methylated cytosines in the striatum may reflect the particularly high basal level expression of *Bcl11b* in basal ganglia as indicated by the protein atlas (**Fig. 3c**)^32^.

*CpGs* in the *Bcl11b^+/−^* knockout methylation signature were located near genes that play a role in the unfolded protein response and ER-nucleus signaling pathway (**Fig. 3e; Extended Data Fig. 10d; Table S4**). The *Bcl11b* heterozygous knockout signature overlapped with the set of CpGs that were found to be significant in our LUC and LUC.adjW analysis (98 out of 560, Fisher exact p= 0.005, **Fig. 3f**). The nominally significant overlap suggests that a subset of the LUC-related CpGs may be regulated by *Bcl11b*.

### Transcriptomic studies in *Bcl11b* knockout mice

We used two publicly available transcriptomic data sets that involved perturbations of *BLC11B*. First, striatal gene expression data from a *Bcl11b* over-expression study in immortalized striatal cells ^33^. Second, striatal gene expression data from a *Bcl11b* homozygous null mice at postnatal day 0 ^34^. The analysis indicated that several genes with LUC CpGs would respond to *Bcl11b* perturbation (**Fig. 3g,h**). We found a nominally significant overlap between the transcriptomic signature of the *Bcl11b* KO in striatum and the genes from our LUC and LUC.adjW signature (Fisher Exact P=0.05, **Fig. 3h**). Conversely, the overexpression of *Bcl11b* affected the expression of only five genes from our LUC and LUC.adjW signature (**Fig. 3g**). Specifically, it resulted in the downregulated expression of *Sox5*, *Trib2*, *Nr2f2*, and *Reck*, but upregulation of *Cxxc4* and *Ptprd*.

### Comparison with the transcriptomic profile of *Polg* knockout mice

*CpGs near* mitochondrial polymerase gamma (*POLG)* were among the top hits in our LUC analysis in mammalian skin. The *Polg^D257A/D257A^* mouse line is a known model of progeria with accelerated mitochondrial point mutations and premature aging phenotypes such as hair loss, graying, spinal curvature, testicular atrophy, decreased bone mass, anemia, weight loss, hearing loss, and premature death ^35^. We analyzed a publicly available transcriptomic dataset from the cochlea of 9 month old *Polg^D257A/D257A^* mice ^36^. The transcriptomic signature associated with *Polg^D257A/D257A^* did not overlap significantly with genes from the LUC signature (**Fig. 3i**, Fisher Exact P=0.5). However, the two sets of genes only shared 15 genes in common (**Fig. 3i**). Thus, this analysis suggests that *Polg* does not have a regulatory effect on genes implicated by our LUC analysis. However, the analysis of *Polg* effects should be revisited with other tissue types such as skin and blood, when these data become available.

### Human GWAS and GenAge database

We overlapped genes from the LUC signature with genes found in human genome-wide association studies (GWAS) of various pathologies and conditions^37^. With all due caution, we report that some of the genes from the LUC signature were those highlighted by GWAS to be associated with type II diabetes (e.g., *ADCY5*, *TCF4*, *BCL11A*), stroke (e.g., *CASZ1*), chronic kidney disease (e.g., *BCAS3*), and breast cancer (e.g., *CREB5*, *FOXP1*) (**Extended Data Fig. 11a**).

We also examined the potential overlap between genes associated with LUC signature and those reported in the GenAge database ^38^, which is a curated collection of longevity genes. We found that 55 genes (14%) of those identified by LUC are also present in the GenAge database, but this overlap was not statistically significant (Fisher Exact P=0.2, **Extended Data Fig. 11b**).

### Epigenetic clocks based on the LUC signature

We and others have previously shown that cytosine methylation levels lend themselves to defining highly accurate estimators of chronological age in many species^5–25, 39–41^. These epigenetic clocks are typically based on elastic net regression with respect to all CpGs on the mammalian methylation array that map to the respective species (Methods). Here we pursued a slightly different strategy to build epigenetic clocks. First, we only used the subset of CpGs that were found to be significant in our LUC (or LUC.adjW) analysis. Second, we used a ridge regression model (as opposed to an elastic net regression model) to ensure that all CpGs were kept in the predictive model. We fit different regression models based on different selections of tissues. Since these clocks are based on LUC, we refer to them as the LUC clocks. Our pan-tissue LUC clock was trained on 1468 samples from 10 different tissue types and is expected to apply to most mouse tissues (**Table S5**). By contrast, tissue-specific LUC clocks were trained on data derived from a single tissue type such as liver (**Table S5**). The underlying multivariate regression model can be used to predict age. The predicted value is often referred to as DNA methylation (DNAm) age or epigenetic age. An application to independent test data reveals that the resulting Uber epigenetic clocks are highly accurate predictors of chronological age (Pearson correlation between the DNAm age estimate and age ranges from r=0.92 to 0.99, **Fig. 4**). To avoid the confounding effect of age, we defined an age-adjusted measure of DNAm age, referred to as epigenetic age acceleration, by regressing DNAm Age on chronological age and forming residuals.

### LUC clock analysis of aging interventions in mice

Growth hormone knock-down mouse lines have been shown to exhibit a lower epigenetic age^42^. Age analyses using the LUC clocks replicated these findings in several tissues, including kidney (p=3e-6), and liver (p=7e-8) (**Fig. 5a**).

Another age intervention is caloric restriction, which has also been shown to be associated with delayed epigenetic aging^42–44^. Once again, analyses of the methylation profiles by our LUC clock replicated these results in livers of calorie-restricted mice (p=1e-10). Conversely, a high-fat diet, which has been shown to increase the epigenetic age of murine liver^45, 46^, was also demonstrated to do so by the LUC clocks (p=0.0008, **Fig. 5a**).

We applied the LUC clocks to the striatum of *Bcl11b* knockout mice. As expected, *Bcl11b* knockout caused an age acceleration in the striatum of these mice (panTissue clock p=5e-5; brain clock p= 0.001) (**Fig. 5b**), which is another piece of indirect evidence supporting the likelihood that *Bcl11b* may regulate LUC signatures.

## DISCUSSION

Here we describe a novel approach for analyzing the methylation data from the Mammalian Methylation Consortium that hones in on divergent aging patterns between long and short-lived species. To quantify this divergence in aging patterns, we introduced an integrative correlation coefficient referred to as LUC. We distinguish two types of LUC analyses: one that ignores differences in body size, and the other that accounts for it. The latter analysis, LUC.adjW, lends itself to finding CpGs and their associated genes that may play a protective role in long-lived species with a relatively small body size. Previous studies have looked at aging-associated DNA methylation dynamics as a molecular readout of lifespan variation among mammalian species^47^. The current study has a different emphasis: it aims to select CpGs with strongly divergent aging patterns in long and short-lived species. While the LUC approach is biologically compelling, it has a statistical limitation: it requires large sample sizes. Our permutation test analysis provides insights into the significance levels that are needed to protect against false positives.

Our LUC results are clearly dependent on tissue type. The most significant LUC CpGs in blood are largely different from those in skin. The poor agreement of LUC results between tissue types is likely due to multiple reasons. First, each tissue type involved a different set of species. For example, we analyzed over 750 blood samples from dogs but did not have any skin samples from this species. Second, the overlap may also reflect biological differences. Genes and underlying CpGs that are associated with an enhanced immune system (i.e., blood) in long-lived species may differ from those that play a protective role in skin. This tissue specificity may indeed be relevant with regards to *POLG*, which was identified by the LUC EWAS in skin, but was not corroborated by transcriptome data from the cochlea of *Polg* mutant mice. Therefore, similar analysis in skin of *Polg* mutant mice ^36, 48^ would be necessary to resolve this question.

Our analysis of blood implicates *BCL11B* which was significant in both LUC and LUC.adjW analyses. B cell leukemia 11 b (*BCL11B*) is a zinc finger protein with a wide range of functions, including the development of the brain, immune system, and cardiac system^49^. This gene is also implicated in several human diseases including, but not limited to, Huntington disease^50^, Alzheimer’s diseases^50^, HIV^51^, and T-cell malignancies^52^. *BCL11B* plays an important role in adult neurogenesis^53^, but is less studied in the context of lifespan disparities in mammals.

We investigated the effect of *Bcl11b^−/+^* knockout on brain methylation levels and transcriptomic changes. Strikingly, we find that *Bcl11b* knockout affected both DNA methylation and mRNA expression of LUC genes. Our current study does not inform us about the potential role of *Bcl11b* in the aging processes during adulthood since the observed patterns could be attributed to developmental defects. Future studies should include conditional *Bcl11b* knockout or knockdown after adulthood to distinguish between the developmental and aging effects of this protein.

We used LUC-related CpGs to develop novel epigenetic age estimators for mice. These LUC clocks do not outperform existing mouse clocks in terms of accuracy of chronological age. Rather, these clocks are meant to address a more elusive aim: the measurement of biological age that correlates with lifespan. The LUC clocks are based on the premise that LUC-related CpGs, being also linked to lifespan, are more informative of biological age than purely age-related CpGs. However, this hypothesis requires validation. The tools to carry out such studies, namely epigenetic clocks, epigenetic predictors of maximum lifespan, and LUC clocks are available to address this hypothesis. Here we demonstrate that LUC clocks show the expected behavior in benchmark anti-aging (caloric restriction, GHRKO) and pro-aging interventions (high-fat diet). Furthermore, the LUC clocks also revealed that *Bcl11b* KO mice undergo accelerated epigenetic aging in the striatum (**Fig. 5c**). We are characterizing other genetic and non-genetic interventions that perturb the LUC clocks, and these will feature in a separate report that will uncover the biological processes that are regulated by the LUC cytosines and their associated genes.

## Supporting information

LUC clock (v1) training set and coefficients

Mammalian Methylation Consortium

A workflow of using LUC version 1 clocks in mice

## Acknowledgment

This work was mainly supported by the Paul G. Allen Frontiers Group (SH) and a grant by Open Philanthropy (SH). The Yang lab (N.W. and X.W.Y.) was supported by NINDS/NIH grants (R01NS113612) and CHDI Foundation, Inc.

## Data Availability

The DNAm data will be made publicly available as part of the data release from the Mammalian Methylation Consortium. Genome annotations of these CpGs can be found on Github https://github.com/shorvath/MammalianMethylationConsortium.

## METHODS

### DNA methylation data

The data used in this study included a total of 8,663 samples from blood, cerebral cortex, liver, muscle, and skin that are collected from 82 mammalian species. These included species had >4 samples at different ages that covered at least 20% of the lifespan. Sample collection and ethical approval for each mammalian species is described in separate individual papers ^5–26, 41^. The species level characteristics such as maximum lifespan, average weight, and age at sexual maturity were chosen from anAge database ^54^. All DNA samples were analyzed by the novel custom-designed mammalian methylation array, which profiles around 36k conserved CpGs across several mammals ^28^. Following data collection, the SeSaMe normalization method was used to define beta values for each probe ^55^. The analysis was limited to 14,705 highly conserved CpGs in eutherian species (description in ^26^).

### *Bcl11b* heterozygous KO mice

*Bcl11b* heterozygous KO mice (Strain name: C57BL/6N-*Bcl11b*^tm1.1(KOMP)Vlcg/JMmucdwere^) (Valenzuela 2003, PMID: 12730667) were obtained from Mutant Mouse Regional Resource & Research Center (MMRRC).

The mouse methylation array study was conducted at UCLA. Animals were housed in standard mouse cages under conventional laboratory conditions, with constant temperature and humidity, 12 h/12 h light/dark cycle (7.00 a.m./7.00 p.m.), and food and water ad libitum. All animal studies were carried out in strict accordance with National Institutes of Health guidelines and approved by the UCLA Institutional Animal Care and Use Committees. C57BL6J mice were used as controls in this experiment. The brain samples were collected at two ages: 0.16 and 0.5 years. A total of 4 animals/group per sex (total of 16 per group) were sacrificed at each age. The EWAS analysis was done by combining both groups and using chronological age as a co-variate in the multivariate regression model.

### Growth hormone receptor KO mice

GHRKO mice (GHR −/− dwarfs) from the University of Michigan (Richard A. Miller). Full-body growth hormone receptor knockout (GHR-KO) ^56^. Sample size: GHRKO, 12 kidney, 11 liver; C57BL6J, 10 kidney, 10 liver. Ages: 0.6-0.7 years.

### Caloric restriction

These data will be described in another publication by Acosta-Rodríguez V and J. Takahashi (submitted).

### Transcriptome datasets

DNAm-mRNA analysis was done in 57 horse transcriptome data generated from 29 different horse tissues ^31^. For humans, we leveraged gene expression studies from 1) GTEx project, which represents multiple tissues and brain regions of 651 men and women, aged 20-80 yr ^57, 58^, 2) three gene expression data studied in ^59^ (GEO datasets from studies ^36, 60, 61^) and 3) the summary data across three studies in ^62^. Mouse datasets included single cell RNA seq data from 23 mouse tissues across the lifespan in the Tabula Muris Consortium ^63^, and microarray data from ^33, 34, 64^. GTEx data: 04/13/2020 and/or dbGaP accession number phs000424.v8.p2 on 04/13/2020.

### Identification of Mappable CpGs for eutherians

Mammalian methylation array contains up to 36k CpGs with high conservation in mammals. However, several of these CpGs only have partial conservation in mammals. Thus, we used stringent criteria to select the CpGs that can be aligned to 11 representative species from different phylogenetic orders ^28^. These species included Human (hg19), mouse (mm10), Vervet monkey (ChlSab1.1.100), Rhesus macaque (Mmul_10.100), Cattle (ARS-UCD1.2), Cat (Felis_catus_9.0.100), Dog (CanFam3.1), Elephant (loxAfr3.100), Bat (Rhinolophus_ferrumequinum.HLrhiFer5), Killer whale (GCF_000331955.2_Oorc_1.1), and Opossum (Monodelphis_domestica.ASM229v1.100). These species are selected based on a large sample size in our data, a relatively high genome quality, and also representation from different phylogenetic orders. This set of probes was further filtered by calibration data generated from the array’s performance on human, mouse, and rat synthetic DNA at different methylation levels (from 0-100% methylated) ^28^. Only the 14,705 probes with a linear correlation of >=0.8 in all three species calibration data were kept as a mappable probe in eutherians.

### Lifespan Uber Correlation analysis

This study introduced “Uber correlation” as a new measure to identify the conserved CpGs with differential aging rates that relate to mammalian lifespan. This analysis comprises of two stages. 1) the CpG-level correlations with age within each species were extracted. 2) the DNAm aging correlation values for each CpGs were correlated with log maximum lifespan of mammalian species. We define the “Lifespan Uber Correlation” (LUC) by correlating the within species aging pattern (defined as Pearson correlation between CpG and chronological age) with maximum lifespan (Fig. 1b). Maximum lifespan was transformed using a log transformation (base e). For each CpG and tissue type, the LUC is implemented in two steps. First, one calculates the Pearson correlation between the CpG and chronological age for all tissue samples from a given species. This step results at a number, F(Species)=cor(CpG,Age), that depends on species (and tissue type). The analysis is restricted to samples of a given tissue type (e.g., blood) to avoid confounding by tissue type. Second, F(Species) is correlated with log transformed maximum lifespan across all available species.

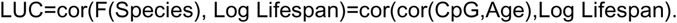

We used the Pearson correlation test implemented in the R function “standardScreeningNumericTrait” function from the “WGCNA” R package ^65^.

For body size independent analysis, the log maximum lifepan of the 82 mammalian species in this study were regressed on log average adult weight of these animals. The residuals of this regression were used as the outcome of the LUC.adjW analysis.

### Permutation analysis

As a sensitivity analysis, the chronological ages of the samples of each species were permuted. This permutation effectively removed the signal between aging pattern and maximum lifespan. The permutation was repeated for 1000 times and the most significant p value, and also maximum correlation value were stored. These permutation test statistics allow us to characterize the distribution of LUC values under the null hypothesis.

### Meta-analysis of transcriptional changes

We aimed to compare the direction of gene expression changes in our target genes in peripheral and brain tissues of mice and humans. In GTEx, the transcription-age association was tested by a mixed-effects model; fixed effects: sex, age; random effect: subjects; outcome: log2 TPM of the genes. For other studies, the summary statistics of the age related gene-expression were available. Stouffer meta-analysis was used to combine the z scores for each tissue with more than one available study. We then extracted the median Z score of the gene expression changes for all the peripheral or brain tissues to do a directional comparison between mice and humans.

### Differential expression analysis

The expression and sample characteristics data were downloaded from NCBI GEO using the “GEOquery” package. Differential gene expression analysis for each experiment was done using linear modeling (Limma package in R).

### Gene ontology enrichment

The genomic region level enrichment was performed using GREAT analysis ^30^, which used 14,705 eutherian CpGs as the background. The analysis used human hg19 annotations, a 50kb window for extending the gene regulatory domain, and default settings for the other options. The biological processes were reduced to parent ontology terms using the “rrvgo” package. The Human GWAS enrichment was done by a hypergeometric overlap analysis between the identified gene regions with the gene sets from 97 published large scale GWAS of various phenotypes. For each GWAS result, we used the MAGENTA software to calculate an overall GWAS P-value per gene, which is based on the most significant SNP association P-value within the gene boundary (+/− 50 kb) adjusted for gene size, number of SNPs per kb, and other potential confounders ^66^.

## Extended data

**Extended Data Fig. 1.**
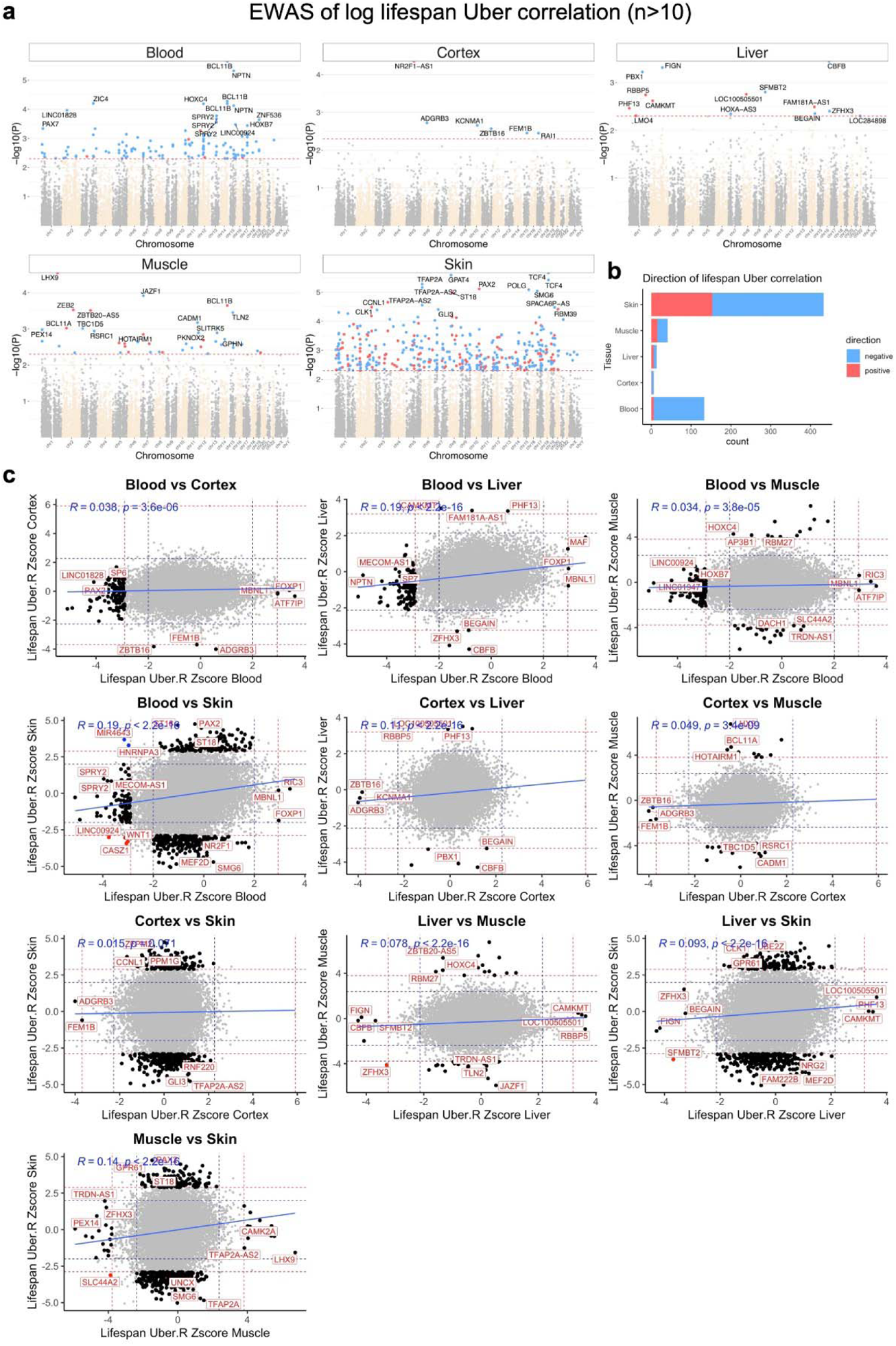
Tissue specific CpGs with differential aging rates related to the maximum lifespan of mammals. **a,** Manhattan plots of the EWAS of differential aging correlation by lifespan in 14,705 conserved CpGs in Eutherians. The coordinates are based on the alignment to Human hg38 genome. The direction of assocoation is highlighted by red (positive correlation with lifespan) and blue (negative correlation with lifespan) at p<0.005 (red dotted line) significance. Only the species with sample size of >10 (range 11 to 1111) and a 20% coverage of lifespan were included in this analysis. Number of species per tissue: Blood, 30 species; Cerebral cortex, 9 species; liver, 21 species; muscle, 12 species; and Skin, 34 species. **b**, stacked bar presentation of the number of significant CpGs per tissues with differential aging correlation with maximum lifespan of mammals. **c**, Pairwise sector plots of differential aging correlation by lifespan in different mammalian tissues. Red-dotted line: p<0.005; blue-dotted line: p>0.05; Red dots: shared CpGs; blue dots: the divergent CpGs.

**Extended Data Fig. 2.**
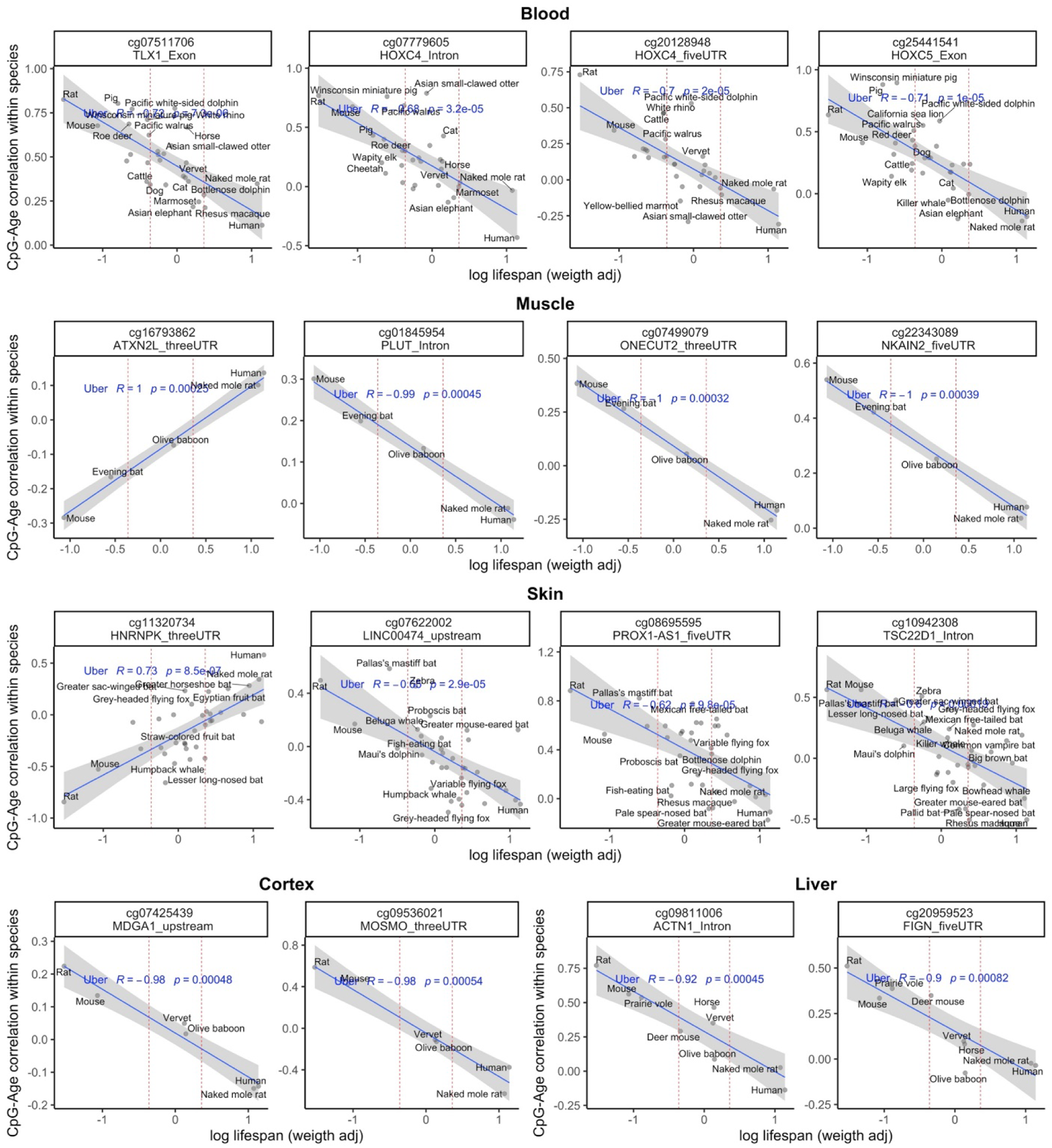
Scatter plots of top CpGs with differential aging correlations associated with mammalian lifespan in different tissues. Pearson correlation coefficients and pvalues are indicated in each figure. The reported gene regions are based on the alignment to Human hg38 genome.

**Extended Data Fig. 3.**
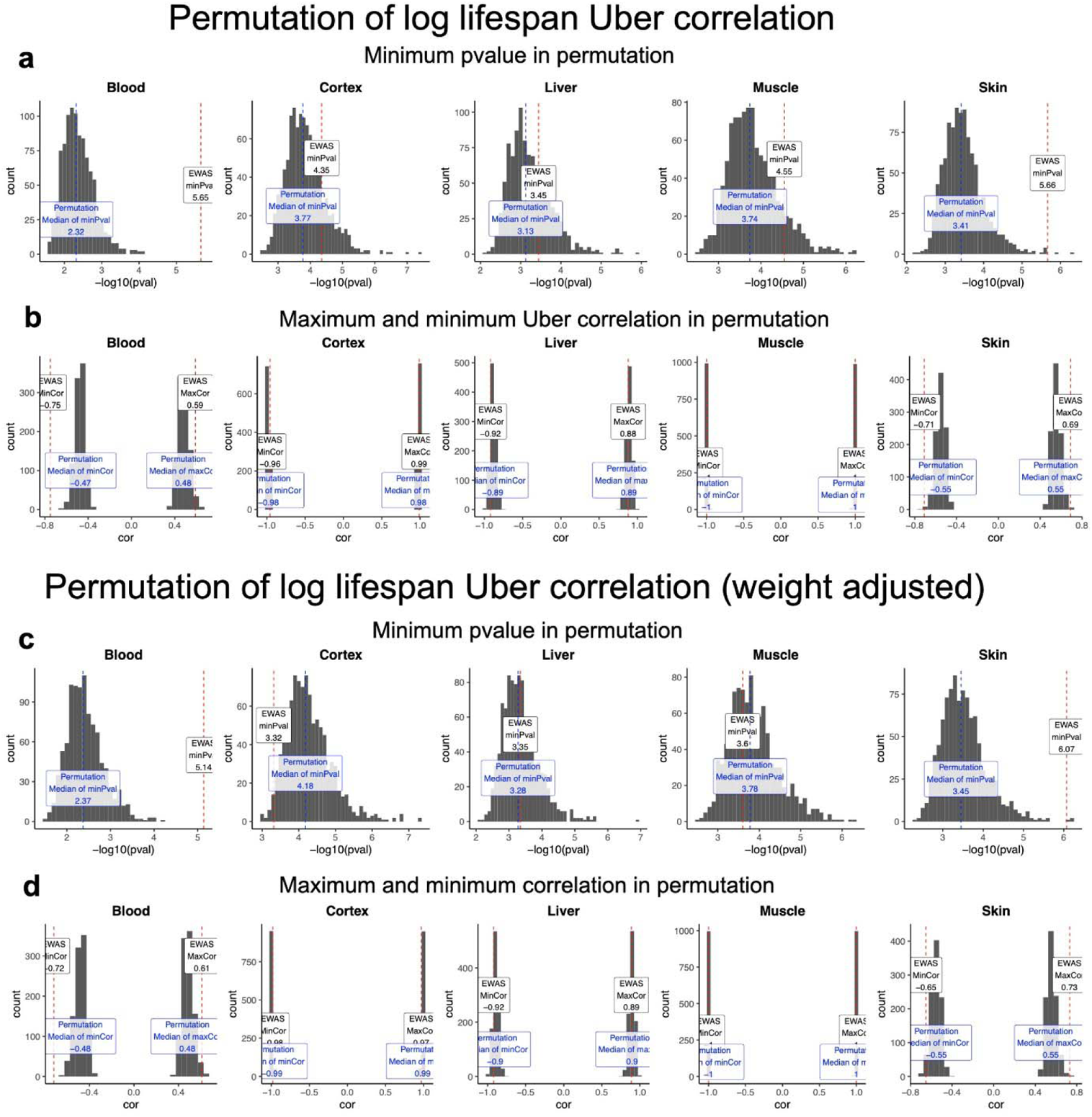
Permutation analysis of the EWAS of differential aging analysis by lifespan or lifespan residuals. Histograms of the minimum pvalues in 1,000 permutation of EWAS of differential aging analysis by lifespan (**a**), or lifespan residuals (**c**). Histograms of the maximum correlation in 1,000 permutation of EWAS of differential aging analysis by lifespan (**b**), or lifespan residuals (**d**). The red dashed lines indicate the actual (unpermuted) EWAS results. The blue dashed lines indicate the median results of the permutation analysis for each tissue.

**Extended Data Fig. 4.**
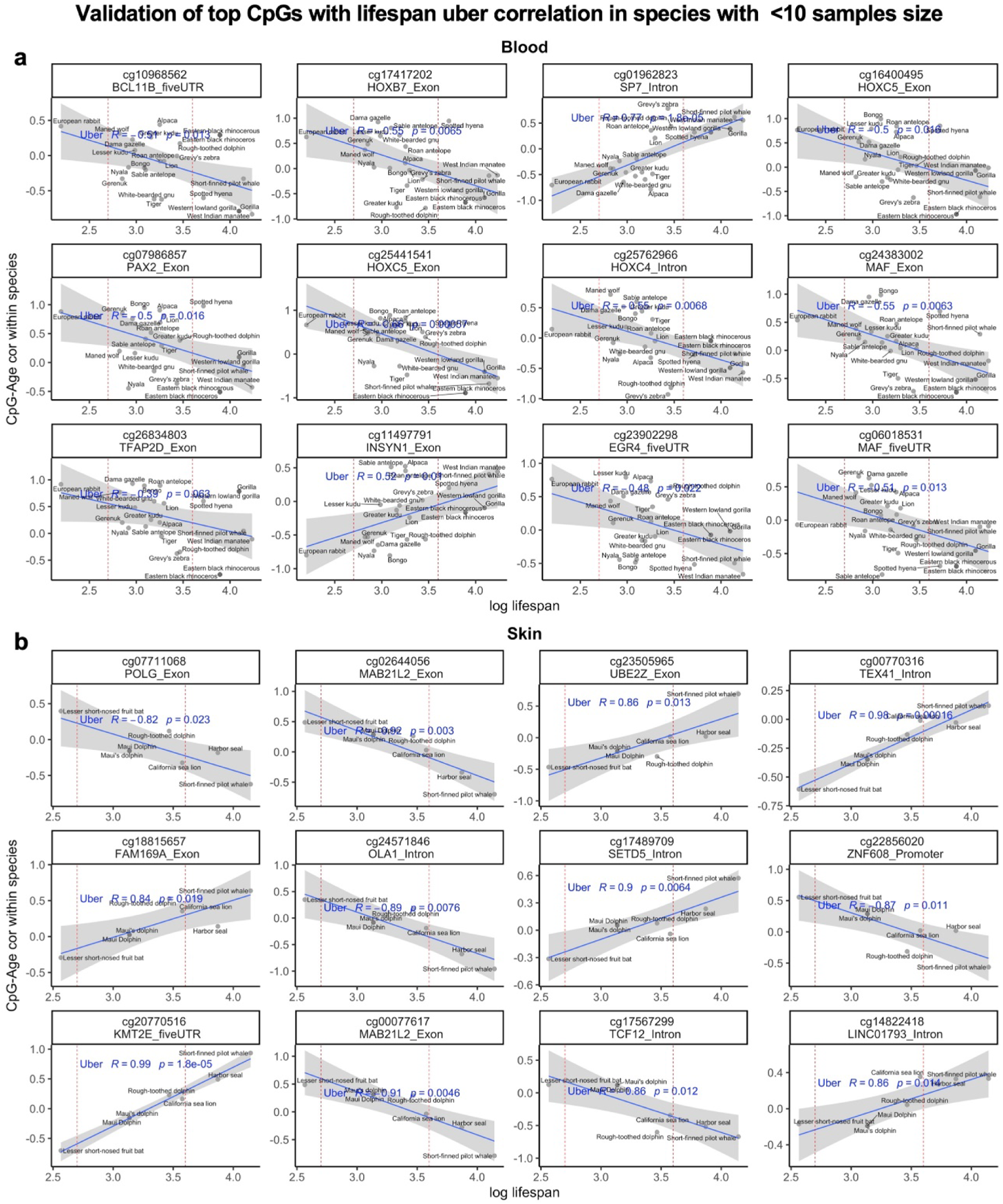
Validation of the top CpGs with differential aging correlation by lifespan in species with less than 10 sample size. The species with small sample size were not included in the main analysis, thus could be used as a validation dataset for the top hits of this study. Number of species per tissues: blood, 23; skin, 6. Scatter plots shows the top EWAS CpGs with corroborated differential aging correlation by lifespan in blood (**a**), and skin (**b**) validation datasets.

**Extended Data Fig. 5.**
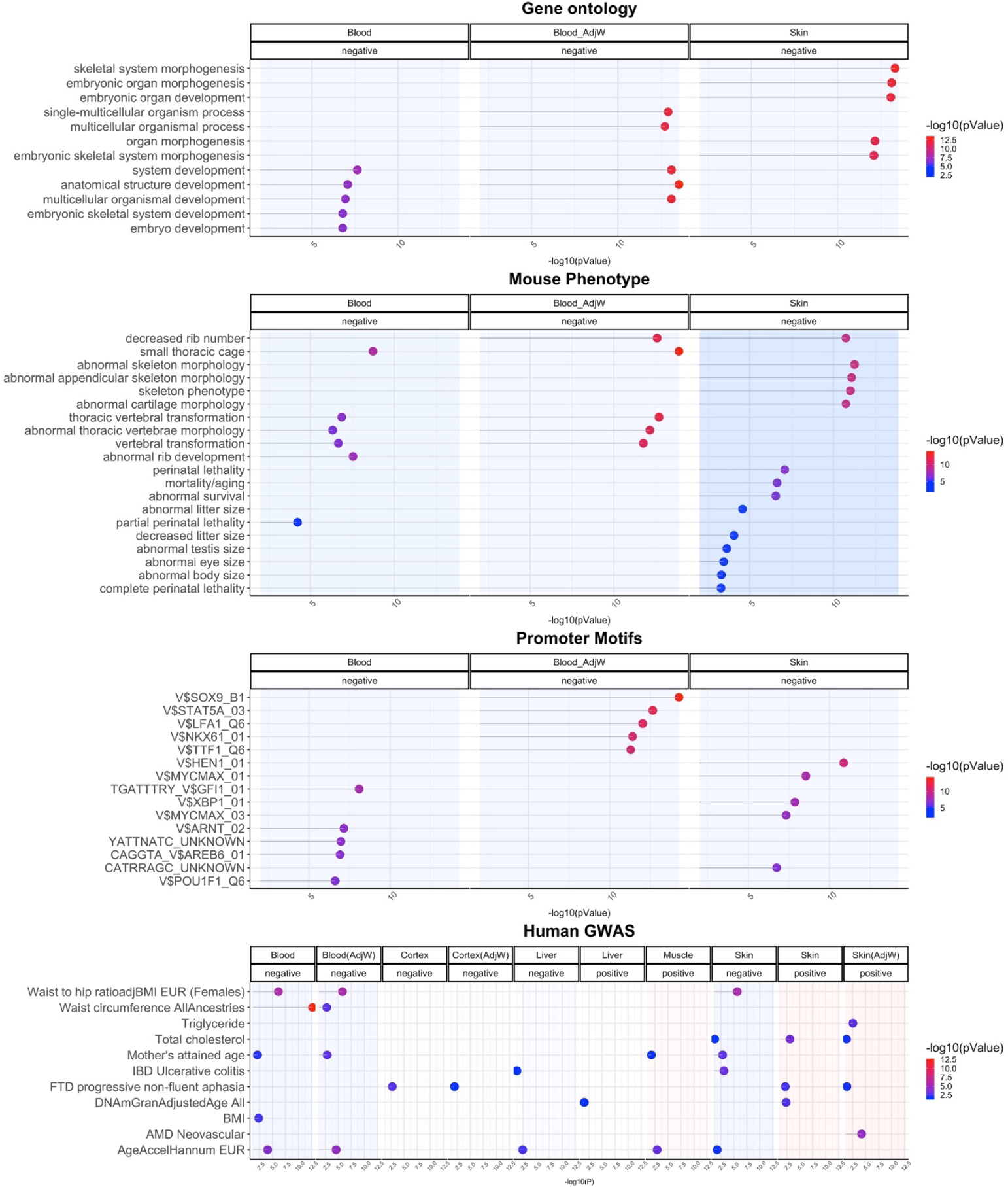
Gene set enrichment analysis of the CpGs with differential aging correlations with mammalian maximum lifespan or lifespan residuals.

**Extended Data Fig. 6.**
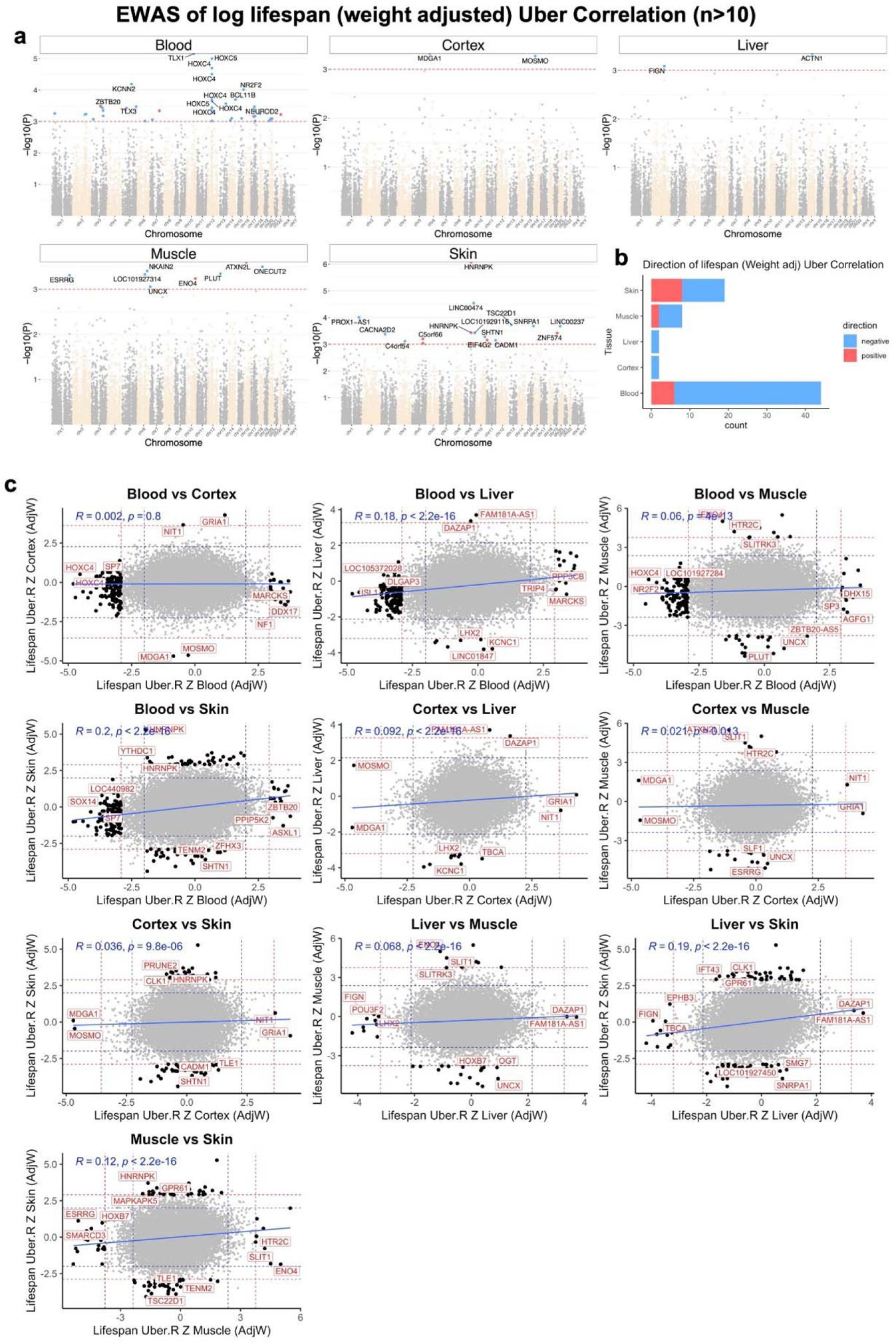
Tissue specific CpGs with differential aging rates related to the residuals of mammalian maximum lifespan adjusted for adult weight of the species. **a,** Manhattan plots of the EWAS of differential aging correlation by lifespan residuals in 14,705 conserved CpGs in Eutherians. The coordinates are based on the alignment to Human hg38 genome. The direction of assocoation is highlighted by red (positive correlation with lifespan) and blue (negative correlation with lifespan) at p<0.005 (red dotted line) significance. Only the species with sample size of >10 (range 11 to 1111) and a 20% coverage of lifespan were included in this analysis. Number of species per tissue: Blood, 30 species; Cerebral cortex, 9 species; liver, 21 species; muscle, 12 species; and Skin, 34 species. **b**, stacked bar presentation of the number of significant CpGs per tissues with differential aging correlation with lifespan residuals in mammals. **c**, Pairwise sector plots of differential aging correlation by lifespan residuals in different mammalian tissues. Red-dotted line: p<0.005; blue-dotted line: p>0.05; Red dots: shared CpGs; blue dots: the divergent CpGs.

**Extended Data Fig. 7.**
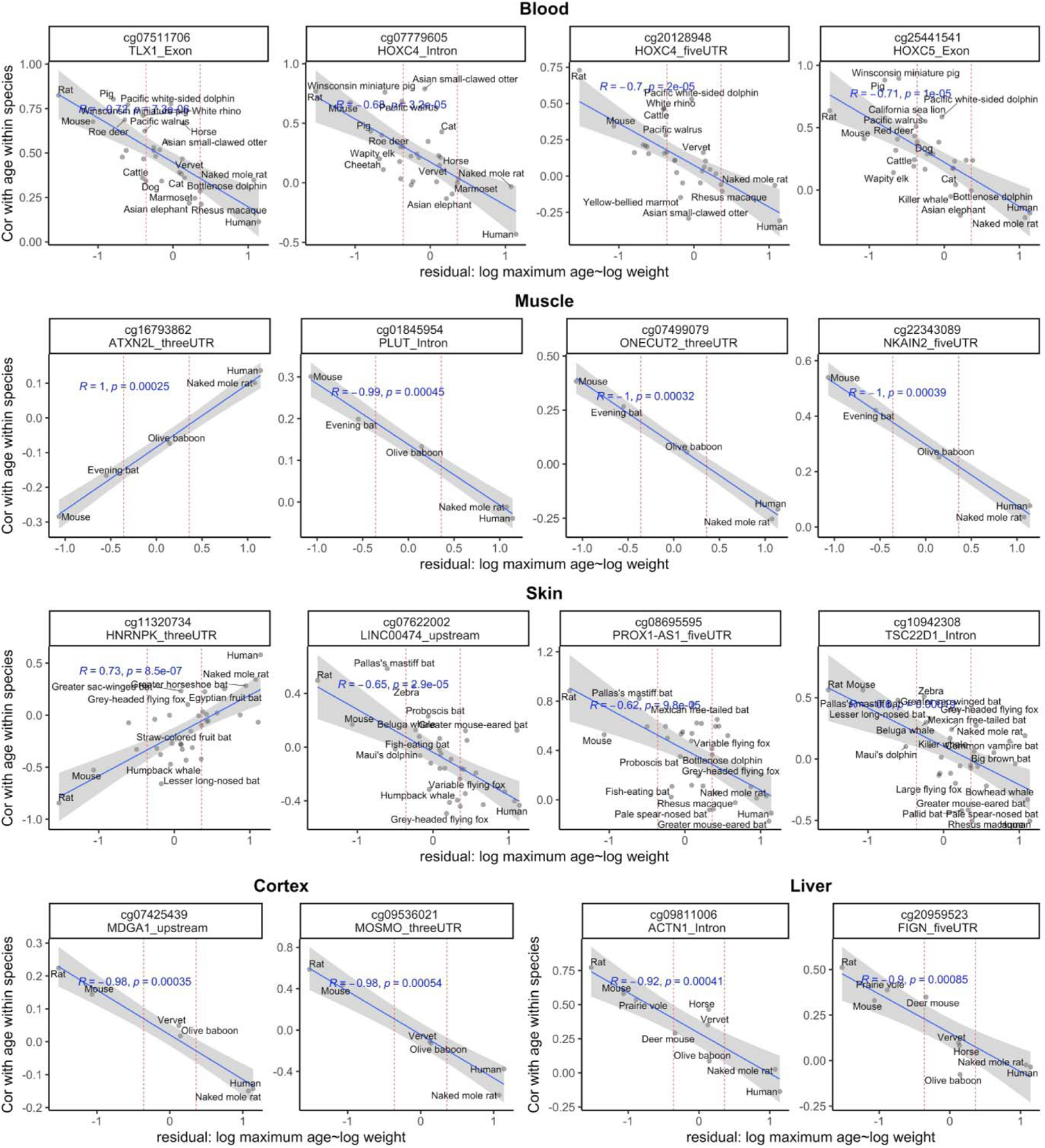
Scatter plots of top CpGs with differential aging correlations assciated with lifespan residuals in different tissues. Pearson correlation coefficients and pvalues are indicated in each figure. The reported gene regions are based on the alignment to Human hg38 genome.

**Extended Data Fig. 8.**
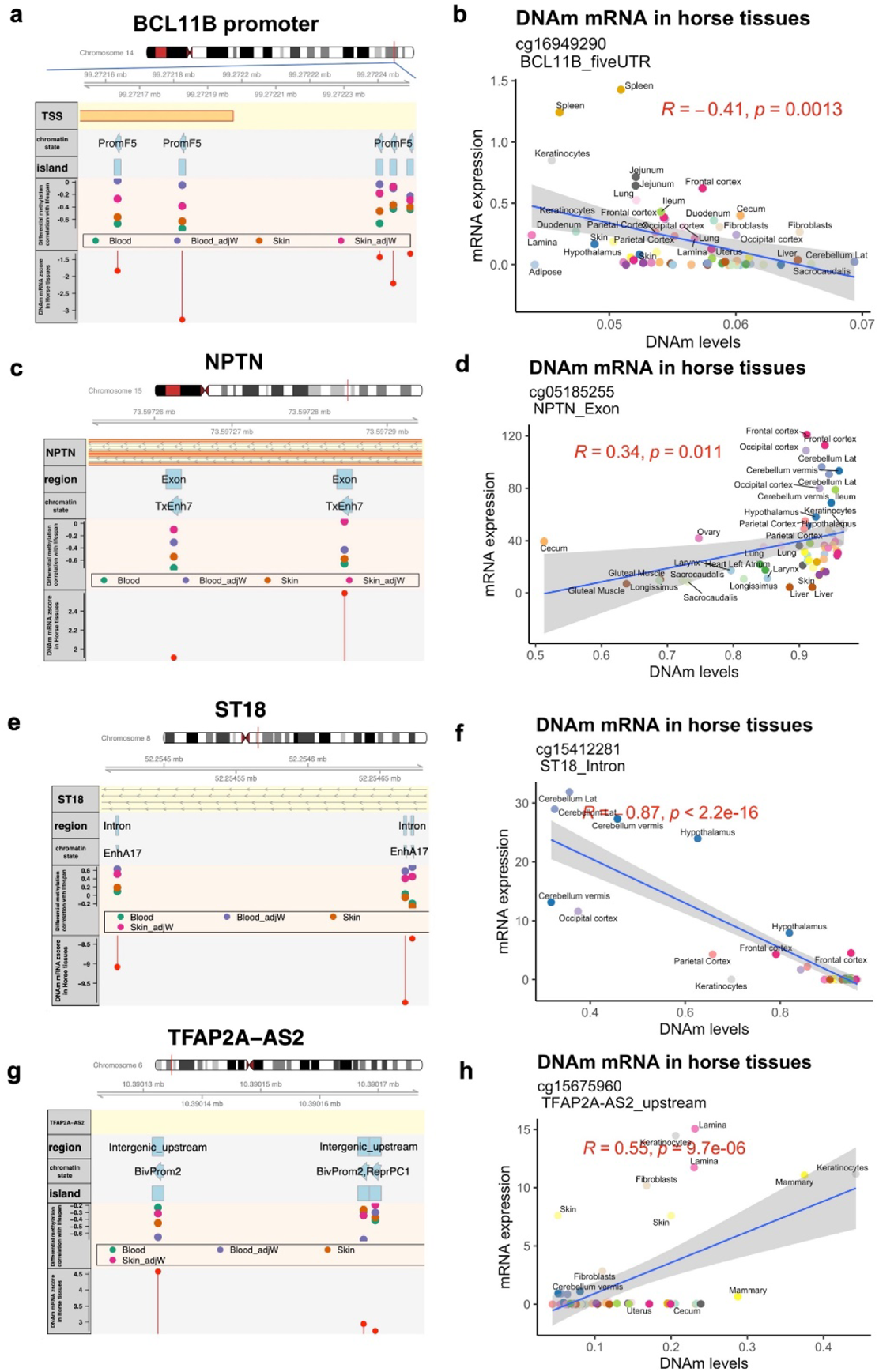
Genome view of the selected CpGs with differential aging correlation by lifespan that relates to gene expression in horse tissues. The genome view is based on human Hg38. For each gene, the CpGs that were significant in at least one analysis were marked in the plot. The genome view plots includes chromosome ideogram, coordinates of the zoomed in region, Ensemble transcript track, gene-regions annotation of highlighted CpGs, chromatin state of highlighted CpGs, CpG island status of the highlighted CpGs, differential aging correlation by mammalian lifespan or lifespan residuals, and the Z score of DNAm-mRNA association in 29 tissues from the two female horses ^31^. Highlighted genes: *BCL11B* (**a**), *NPTN* (**c**), *ST18* (**e)**, and *TFAP2A-AS2* (**g**). The scatter plot of DNAm-mRNA association in horse tissues were illustrated for the selected CpGs in *BCL11B* (**b**), *NPTN* (**d**), *ST18* (**f**), and *TFAP2A-AS2* (h). The plots were created using “Gviz” package in R.

## Age-related mRNA changes in the identified genes in humans and mice

To evaluate whether CpGs with significant LUC are adjacent to genes whose mRNA levels also show a divergent aging pattern between long lived and short lived species, we analyzed several publicly available transcriptomic datasets of human and mouse tissues.

When considering publicly available transcriptomic data from humans and mice, we find opposing aging patterns across tissues from the same species (**Extended Data Fig. 9b,c**). For example, *BCL11* is positively correlated with age in human skeletal muscle, adipose, tibial artery but is negatively correlated with age in human prefrontal cortex, temporal cortex, pons, and other brain regions (**Extended Data Fig. 9b**). As expected, we find opposing aging patterns between species. While BCL11 has a positive correlation with age in human skeletal muscle, it exhibits a significant negative correlation in murine limb muscles. This high divergence of transcriptomic aging patterns can be observed for thousands of genes, i.e. it is not specific to genes that have been implicated in the LUC analysis.

To remove the confounding effect of tissue type and sequencing platform, we carried out meta analysis across tissue type and data set to quantify the relationship between mRNA and age in each species (**Extended Data Fig. 9a**). Our meta analysis statistic was defined as the median Z score of the age correlation test across tissue types (peripheral tissues or brain). The analysis revealed that many genes exhibit transcriptomic aging patterns that differ between the two species.

For example, *CLK1* mRNA increased with age in humans (p=5e-0.3) but it showed the opposite aging pattern in mouse tissues (p=2e-20). Similarly, *HNRNPA3* expression increased weakly with age in human peripheral tissues but decreased very significantly in mouse tissues (p=0.03 in humans, p= 4e-19 in mice). Several genes implicated by the LUC analysis presented with an age related decrease in mRNA in mice but not in humans (*Polg,* p= 1e-10 mice, *NPTN* p=4e-54 in mice, *HNRNPK p=1e-26* mice). The meta analysis results for BCL11B were less striking (nominally significant evidence that *Bcl11b* increases with age in murine brain regions but no significant associations in human brain regions or in peripheral tissues).

**Extended Data Fig. 9.**
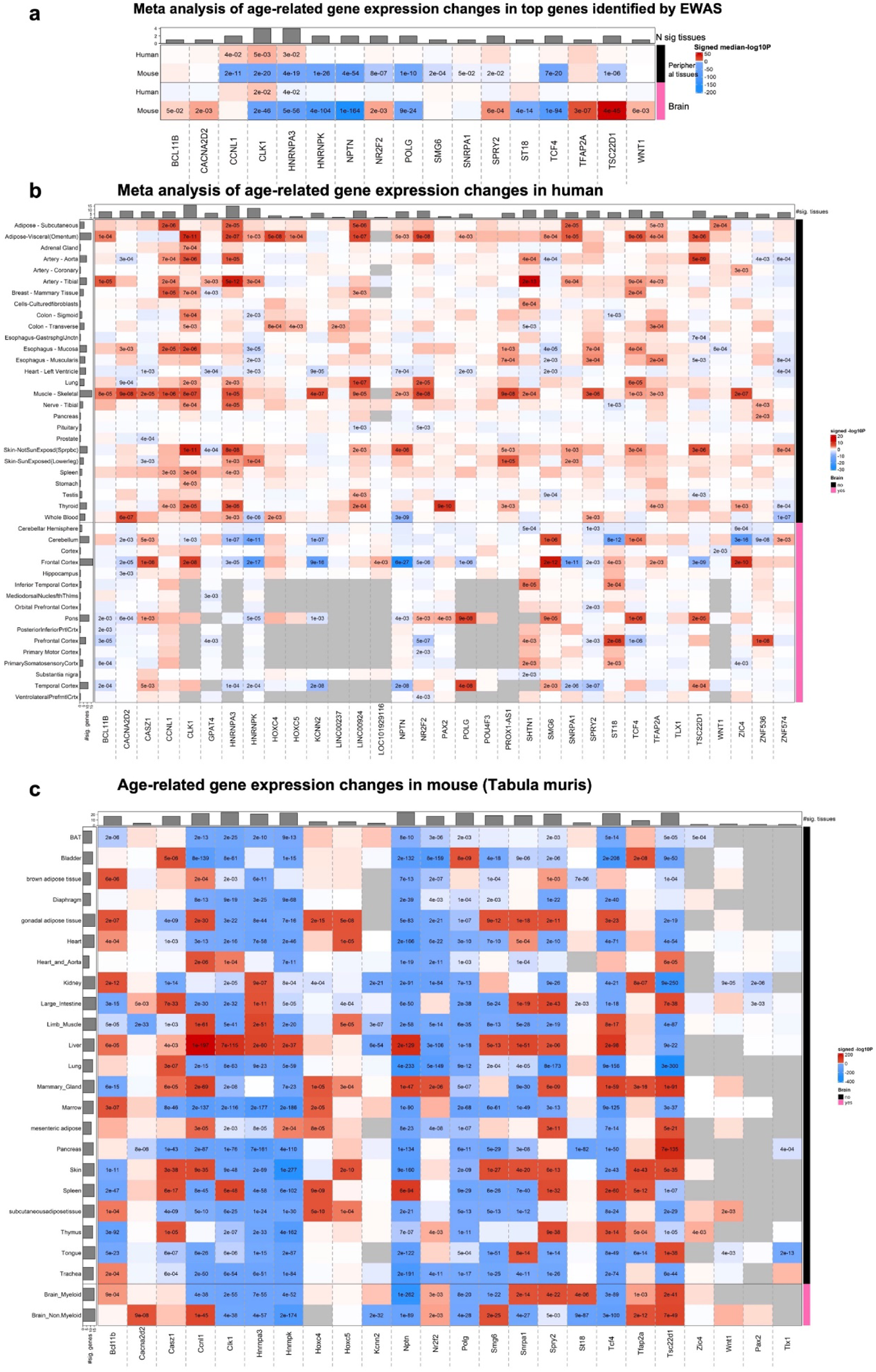
Meta analysis of age related gene expression changes in the top identified genes in different human and mouse tissues. **a,** the meta-analysis of age-related mRNA changes in peripheral and brain tissues of human and mouse. The heatmap represents the median of z scores for different peripheral tissues and brain regions. We used Stouffer meta-analysis z scores if there was more than one dataset available for the tissue. Transcriptome datasets: Human (**b**)^57, 67^, Mouse (**c**) ^64^. Age related mRNA changes was assessed by multivariate model including sex, and batch as co-variates.

**Extended Data Fig. 10.**
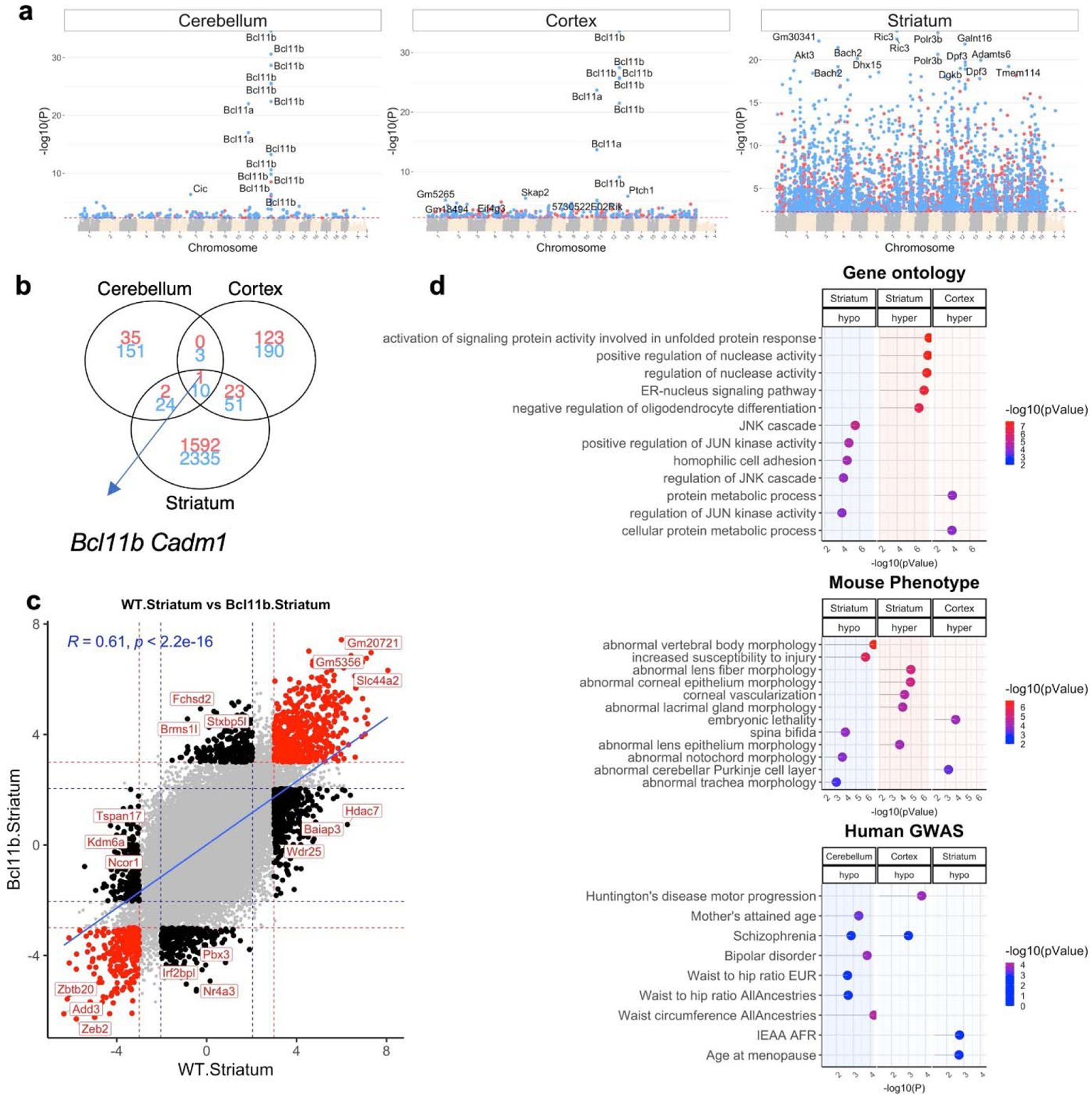
*Bcl11b^−/+^* knock out mainly cause DNAm changes in the striatum. **a,** Manhattan plots of the EWAS of *Bcl11b* heterozygous knockout in the striatum, cerebral cortex and cerebellum of mice. The coordinates are based on the alignment to Mouse mm10 genome. The direction of association is highlighted by red (positive association) and blue (negative association) at p<0.005 (red dotted line, FDR<0.03) significance. Sample size: 16/group, ages, 0.16-0.5 years. The association was done by a multivariate model with age as co-variate. **b,** venn diagram of the significant EWAS results in each brain region. **c,** scatter plot of DNAm aging in Bcl11b^−/+^ vs wildtype mouse lines. Red dots indicate the shared age related CpGs at p<0.005. **d,** gene set enrichment analysis of the genes associated with Bcl11b knockout.

**Extended Data Fig. 11.**
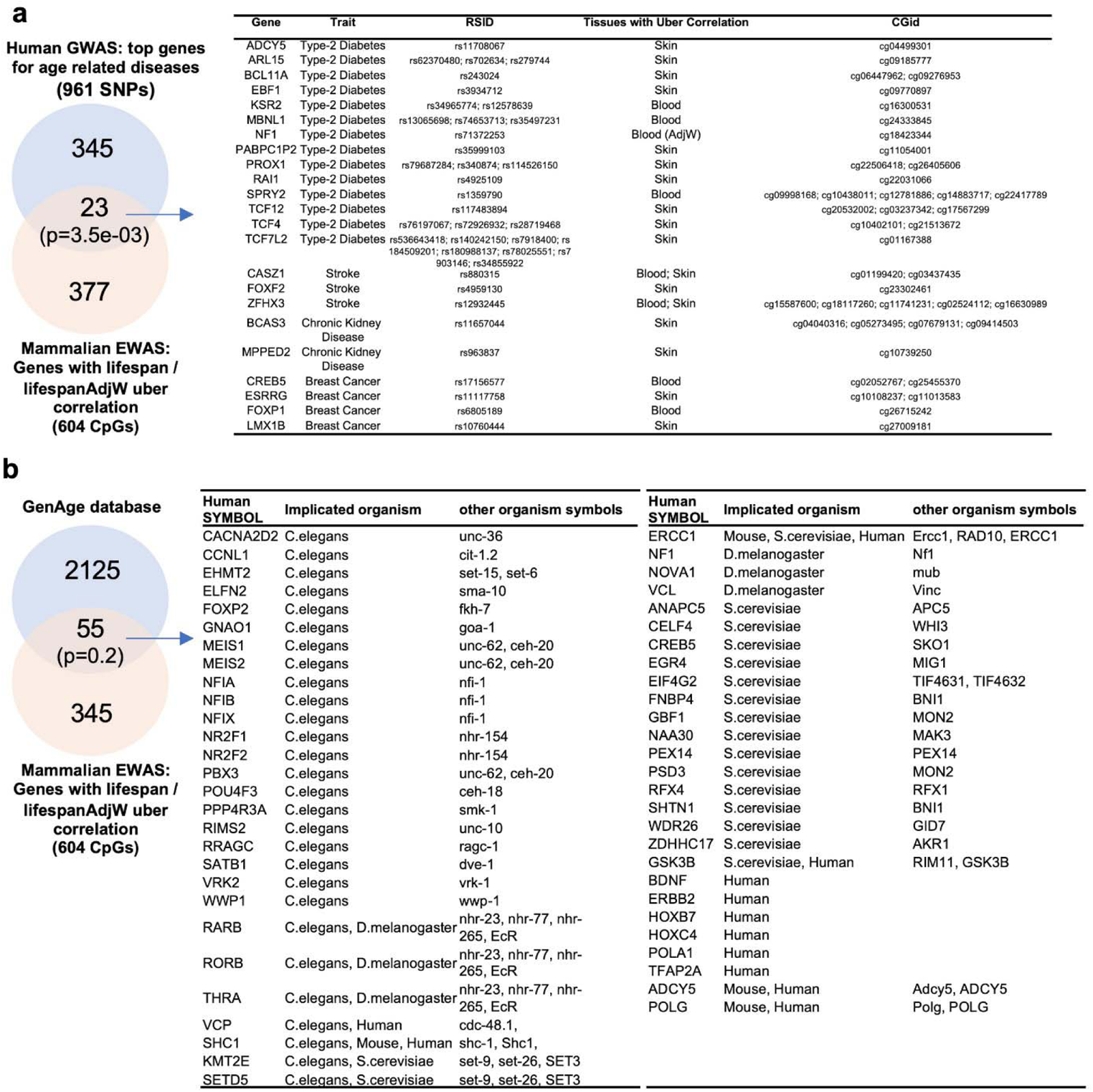
Genes from the LUC or LUC.adjW signature versus GWAS and GenAge. **a**, Mammalian LUC (and LUC.adjW) exhibit a nominally significant overlap with genes that have been found by GWAS of human chronic diseases ^37^. **b,** The overlap of the mammalian LUC (and LUC.adjW) results with longevity genes reported in GenAge database ^38^. The pvalues in Venn diagram report the Fisher exact test results of the overlap of 400 LUC (or LUC.adjW) genes with each dataset. The background was limited to the genes that are covered in Mammalian Methylation Array.

