## Supplementary material for "Divergent age-related methylation patterns in long and short-lived mammals": Mammalian Methylation Consortium

(Alphabetic order)

Ablaeva, J.[1]; Acosta-Rodríguez, V.A.[2]; Adams, D.M.[3]; Alagaili, A.N.[4], [4]; Almunia, J.[5], [5]; Aloysius, A.[6], [6]; Amor, N.M.S.[4]; Ardehali, R.[7]; Arneson, A.[8], [37]; Baker, C.S.[9]; Banks, G.[10]; Belov, K.[11], [11]; Bennett, N.C.[12], [12]; Black, P.[13]; Blumstein, D.T.[14], [115]; Bors, E.K.[9]; Breeze, C.E.[15], [15]; Brooke, R.T.[16], [16]; Brown, J.L.[17]; Carter, G.[18]; Caulton, A.[19], [116]; Cavin, J.M.[20]; Chakrabarti, L.[21], [21]; Chatzistamou, I.[22]; Chavez, A.S.[23], [117]; Chen, H.[24]; Cheng, K.[25]; Chiavellini, P.[26]; Choi, O.W.[27]; Clarke, S.[19], [19]; Cook, J.A.[28], [28]; Cooper, L.N.[29]; Cossette, M.L.[30], [30]; Day, J.[31]; DeYoung, J.[27]; Dirocco, S.[32]; Dold, C.[33]; Dunnum, J.L.[28], [28]; Ehmke, E.E.[34]; Emmons, C.K.[35]; Emmrich, S.[1]; Erbay, E.[36], [118]; Erlacher-Reid, C.[32]; Ernst, J.[37], [119]; Faulkes, C.G.[38], [38]; Fei, Z.[39], [39]; Ferguson, S.H.[40], [120]; Finno, C.J.[41], [41]; Flower, J.E.[42]; Gaillard, J.M.[43]; Garde, E.[44]; Gerber, L.[45], [121]; Gladyshev, V.N.[46], [46]; Gorbunova, V.[1], [1]; Goya, R.G.[26], [26]; Grant, M.J.[47], [47]; Green, C.B.[2]; Haghani, A.[48], [48]; Hanson, M.B.[35]; Hart, D.W.[12], [12]; Haulena, M.[49]; Herrick, K.[50]; Hogan, A.N.[51], [51]; Hogg, C.J.[11], [11]; Hore, T.A.[52], [52]; Horvath, S.[48], [122]; Huang, T.[53], [123]; Izpisua Belmonte, J.C.[54]; Jasinska, A.J.[27]; Jones, G.[55]; Jourdain, E.[56]; Kashpur, O.[57]; Katcher, H.[58]; Katsumata, E.[59]; Kaza, V.[60]; Kiaris, H.[61], [124]; Kobor, M.S.[62]; Kordowitzki, P.[63], [125]; Koski, W.R.[64], [64]; Kruetzen, M.[65], [65]; Kwon, S.B.[37], [119]; Larison, B.[14], [126]; Lee, S.G.[46], [46]; Lehmann, M.[26]; Lemaitre, J.F.[43]; Levine, A.J.[66]; Li, C.[68], [127]; Li, C.Z.[39], [39]; Li, X.[67]; Lim, A.R.[48]; Lin, D.T.S.[62]; Lindemann, D.M.[32]; Liphardt, S.W.[69], [69]; Little, T.[70], [70]; Lu, A.T.[48], [48]; Macoretta, N.[1]; Maddox, D.[71]; Matkin, C.O.[72]; Mattison, J.A.[73]; McClure, M.[74]; Mergl, J.[75]; Meudt, J.J.[76]; Miller, R.A.[77], [77]; Montano, G.A.[78]; Mozhui, K.[79], [128]; Munshi-South, J.[80], [80]; Murphy, W.J.[81], [129]; Naderi, A.[61], [61]; Nagy, M.[82]; Narayan, P.[47]; Nathanielsz, P.W.[68], [127]; Nguyen, N.B.[7]; Niehrs, C.[83], [130]; Nyamsuren, B.[84]; O'Brien, J.K.[31], [31]; O'Tierney Ginn, P.[57]; Odom, D.T.[85], [131]; Ophir, A.G.[86]; Osborn, S.[87]; Ostrander, E.A.[51], [51]; Parsons, K.M.[88], [88]; Paul, K.C.[66]; Pellegrini, M.[89]; Peters, K.J.[65], [132]; Petersen, J.L.[90]; Pietersen, D.W.[91], [133]; Pinho, G.M.[14]; Plassais, J.[51], [51]; Poganik, J.[46], [46]; Prado, N.A.[17], [134]; Raj, K.[92], [92]; Reddy, P.[93]; Rey, B.[43]; Ritz, B.R.[94], [66]; Robbins, J.[95], [95]; Robeck, T.R.[78], [78]; Rodriguez, M.[96]; Russell, J.[87], [50]; Rydkina, E.[1]; Sailer, L.L.[86]; Salmon, A.B.[97]; Sanghavi, A.[58]; Schachtschneider, K.M.[98], [135]; Schmitt, D.[99]; Schmitt, T.[100]; Schomacher, L.[83]; Schook, L.B.[98], [136]; Sears, K.E.[14], [14]; Seifert, A.W.[6], [6]; Seluanov, A.[1], [1]; Shafer, A.B.A.[101], [101]; Shindyapina, A.V.[46], [46]; Simmons, M.[34]; Singh, K.[102]; Sinha, I.[14]; Slone, J.[53]; Snell, R.G.[47]; Soltanmohammadi, E.[61]; Spangler, M.L.[103]; Spriggs, M.[13]; Staggs, L.[32]; Stedman, N.[13], [13]; Steinman, K.J.[104], [104]; Stewart, D.T.[105]; Sugrue, V.J.[52]; Szladovits, B.[106], [106]; Takahashi, J.S.[2], [137]; Takasugi, M.[1]; Teeling, E.C.[107]; Thompson, M.J.[89], [89]; Van Bonn, B.[108]; Vernes, S.C.[109], [138]; Villar, D.[110], [110]; Vinters, H.V.[111]; Vu, H.[8]; Wallingford, M.C.[57], [139]; Wang, N.[112], [140]; Wilkinson, G.S.[3], [3]; Williams, R.W.[113]; Yan, Q.[39], [39]; Yang, X.W.[112], [140]; Yao, M.[39]; Young, B.G.[40], [40]; Zhang, B.[46], [46]; Zhang, J.[48], [48]; Zhang, Z.[1]; Zhao, P.[7], [141]; Zhao, Y.[1]; Zhou, W.[114], [142]; Zoller, J.A.[39], [39]

### Affiliations


[1], Depts. of Biology and Medicine, University of Rochester, Rochester, NY, USA;
[2], Department of Neuroscience, Peter O'Donnell Jr. Brain Institute, University of Texas Southwestern Medical Center, Dallas, TX, US;
[3], Dept. of Biology, University of Maryland, College Park, USA;
[4], KSU Mammals Research Chair, Department of Zoology, College of Science, King Saud University, Riyadh, Saudi Arabia;
[5], Loro Parque Fundacion, Avenida Loro Parque, Puerto de la Cruz, Tenerife, Spain;
[6], Department of Biology, University of Kentucky, Lexington, KY, USA;
[7], Division of Cardiology, Dept. of Internal Medicine, David Geffen School of Medicine, University of California, Los Angeles, Los Angeles, CA, USA;
[8], Bioinformatics Interdepartmental Program, University of California, Los Angeles, CA, USA;
[9], Marine Mammal Institute, Oregon State University, Newport, OR, USA;
[10], Mammalian Genetics Unit, MRC Harwell Institute, Harwell Science and Innovation Campus, Oxfordshire, UK;
[11], School of Life and Environmental Sciences, The University of Sydney, Sydney, New South Wales, Australia;
[12], Department of Zoology and Entomology, University of Pretoria, Private Bag X[20], Hatfield, [0028], South Africa;
[13], Busch Gardens Tampa, Tampa, Florida, USA;
[14], Dept. of Ecology and Evolutionary Biology, UCLA, Los Angeles, CA, USA;
[15], Altius Institute for Biomedical Sciences, Seattle, WA, USA;
[16], Epigenetic Clock Development Foundation, Los Angeles, CA, USA;
[17], Center for Species Survival, Smithsonian Conservation Biology Institute, Front Royal, VA, USA;
[18], Dept. of Evolution, Ecology and Organismal Biology, The Ohio State University, USA;
[19], AgResearch, Invermay Agricultural Centre, Mosgiel, Otago, New Zealand;
[20], Gulf World Marine Park - Dolphin Company, Panama City Beach, FL, USA;
[21], School of Veterinary Medicine and Science, University of Nottingham, UK.;
[22], Department of Pathology, Microbiology & Immunology, School of Medicine, University of South Carolina, SC, USA;
[23], Department of Evolution, Ecology and Organismal Biology, The Ohio State University, Columbus, OH, United States;
[24], Dept. of Pharmacology, Addiction Science and Toxicology, The University of Tennessee Health Science Center, Memphis, TN, USA;
[25], Medical Informatics, David Geffen School of Medicine, University of California Los Angeles, Los Angeles, CA, USA;
[26], Biochemistry Research Institute of La Plata, Histology and Pathology, School of Medicine, University of La Plata, La Plata, Argentina;
[27], Center for Neurobehavioral Genetics, Semel Institute for Neuroscience and Human Behavior, Dept. of Psychiatry and Biobehavioral Sciences, David Geffen School of Medicine, University of California Los Angeles, Los Angeles, California, USA;
[28], University of New Mexico, Department of Biology and Museum of Southwestern Biology, Albuquerque, New Mexico, United States of America;
[29], Dept. of Anatomy and Neurobiology, Northeast Ohio Medical University, Rootstown, Ohio, USA;
[30], Department of Environmental & Life Sciences, Trent University, Peterborough Ontario, Canada K[9]J [7]B[8];
[31], Taronga Conservation Society Australia;
[32], SeaWorld Orlando, [7007] SeaWorld Drive, Orlando, FL USA;

[33], Zoological Operations, SeaWorld Parks and Entertainment, [7007] SeaWorld Drive, Orlando, Flo
rida, USA;
[34], Duke Lemur Center, Durham, North Carolina, USA;
[35], Conservation Biology Division, Northwest Fisheries Science Center, National Marine Fisherie
s Service, National Oceanic and Atmospheric Administration, Seattle, Washington, USA;
[36], Dept. of Cardiology, Dept. of Biomedical Sciences, Cedars-
Sinai Medical Center, Los Angeles, CA, USA;
[37], Dept. of Biological Chemistry, University of California, Los Angeles, Los Angeles, California,
USA;
[38], School of Biological and Chemical Sciences, Queen Mary University of London, London, UK;
[39], Dept. of Biostatistics, Fielding School of Public Health, University of California Los Angeles, L
os Angeles, CA, USA;
[40], Fisheries and Oceans Canada, University Crescent, Winnipeg, Canada;
[41], Dept. of Population Health and Reproduction, University of California, Davis School of Veteri
nary Medicine, Davis, CA, USA;
[42], Mystic Aquarium, Mystic, Connecticut, USA;
[43], University of Lyon, CNRS, Laboratoire de Biométrie et Biologie Evolutive, UMR[5558], Villeurba
nne, France;
[44], Greenland Institute of Natural Resources, Nuuk, Greenland;
[45], School of Biological, Earth and Environmental Sciences, University of New South Wales, Syd
ney, Australia.;
[46], Division of Genetics, Dept. of Medicine, Brigham and Women's Hospital, Harvard Medical Sc
hool, Boston, MA, USA;
[47], Applied Translational Genetics Group, School of Biological Sciences, Centre for Brain Resea
rch, The University of Auckland, Auckland, New Zealand;
[48], Dept. of Human Genetics, David Geffen School of Medicine, University of California Los Ang
eles, Los Angeles, CA, USA;
[49], Vancouver Aquarium, Vancouver, Canada;
[50], SeaWorld San Diego, San Diego, CA, USA;
[51], Cancer Genetics and Comparative Genomics Branch, National Human Genome Research In
stitute, National Institutes of Health, Bethesda, MD, USA;
[52], Dept. of Anatomy, University of Otago, Dunedin, New Zealand;
[53], Division of Human Genetics, Department of Pediatrics, University at Buffalo, New York, USA;
[54], Altos Labs, San Diego, CA, USA;
[55], School of Biological Sciences, University of Bristol, Bristol, UK;
[56], Norwegian Orca Survey, Andenes, Norway;
[57], Mother Infant Research Institute, Tufts Medical Center, Boston, MA, USA;
[58], Nugenics Research Pvt Ltd, India;
[59], Kamogawa Sea World, Kamogawa, Chiba, Japan;
[60], Peromyscus Genetic Stock Center, University of South Carolina, SC, USA;
[61], Dept. of Drug Discovery and Biomedical Sciences, College of Pharmacy, University of South
Carolina, SC, USA;
[62], Centre for Molecular Medicine and Therapeutics, BC Children's Hospital Research Institute,
University of British Columbia, Vancouver, Canada;
[63], Institute of Animal Reproduction and Food Research of Polish Academy of Sciences, Olsztyn
, Poland;
[64], LGL Limited, King City, ON, Canada;
[65], Evolutionary Genetics Group, Department of Anthropology, University of Zurich, Zurich, Switz
erland;
[66], Dept. of Neurology, David Geffen School of Medicine, University of California Los Angeles, L
os Angeles, CA, USA;

[67], Technology Center for Genomics and Bioinformatics, Dept. of Pathology and Laboratory Medicine, University of California, Los Angeles, Los Angeles, CA, USA;
[68], Texas Pregnancy and Life-course Health Center, Southwest National Primate Research Center, San Antonio, Texas, USA;
[69], Biology Department, University of New Mexico, Albuquerque, NM, USA;
[70], Institute of Evolutionary Biology, School of Biological Sciences, University of Edinburgh, Edinburgh, UK;
[71], White Oak Conservation Center, Yulee FL, USA;
[72], North Gulf Oceanic Society, Homer, Alaska,USA;
[73], Translational Gerontology Branch, National Institute on Aging Intramural Research Program, National Institutes of Health, USA;
[74], ABS Global, DeForest, WI, USA;
[75], Marineland of Canada, Niagara Falls, Ontario, Canada;
[76], Biomedical and Genomic Research Group, Dept. of Animal and Dairy Sciences, University of Wisconsin Madison, Madison, Wisconsin, US;
[77], Department of Pathology, University of Michigan Geriatrics Center, Ann Arbor, MI, USA;
[78], Zoological Operations, SeaWorld Parks and Entertainment, Orlando, Florida, USA;
[79], Dept. of Preventive Medicine, University of Tennessee Health Science Center, College of Medicine, Memphis, TN, USA;
[80], Louis Calder Center - Biological Field Station, Department of Biological Sciences and Center for Urban Ecology, Fordham University, Armonk, NY USA;
[81], Department of Veterinary Integrative Biosciences, Texas A&M University, College Station, Texas [77843], USA;
[82], Museum fur Naturkunde, Leibniz-Institute for Evolution and Biodiversity Science, Berlin, Germany;
[83], Institute of Molecular Biology gGmbH, Mainz, Germany;
[84], Mammalogical Society of Mongolia, Ulaanbaatar, Mongolia;
[85], Cancer Research UK Cambridge Institute, University of Cambridge, Robinson Way, Cambridge, CB[2] [0]RE, UK;
[86], Dept. of Psychology, Cornell University, Ithaca, NY, USA;
[87], SeaWorld San Antonio, San Antonio, Texas, USA;
[88], Conservation Biology Division, Northwest Fisheries Science Center, National Marine Fisheries Service, National Oceanic and Atmospheric Administration, Seattle, Washington, USA;
[89], Dept. Molecular Cell and Developmental Biology, University of California Los Angeles, Los Angeles, CA, USA;
[90], Dept. of Animal Science, University of Nebraska, Lincoln, NE, USA;
[91], Mammal Research Institute, Department of Zoology and Entomology, University of Pretoria, Private Bag X[20], Hatfield, [0028], South Africa.;
[92], Radiation Effects Dept., Centre for Radiation, Chemical and Environmental Hazards, Public Health England, Chilton, Didcot, UK;
[93], Salk Institute for Biological Studies, La Jolla, CA USA;
[94], Dept. of Epidemiology and Environmental Health Sciences, UCLA Fielding School of Public Health, Los Angeles, CA, USA;
[95], Center for Coastal Studies, Provincetown, MA, USA;
[96], Miami Seaquarium, Miami, Florida, USA;
[97], The Sam and Ann Barshop Institute for Longevity and Aging Studies, and Dept. of Molecular Medicine, UT Health San Antonio, and the Geriatric Research Education and Clinical Center, South Texas Veterans Healthcare System, San Antonio TX, USA;
[98], Dept. of Radiology, University of Illinois at Chicago, Chicago, IL, USA;
[99], College of Agriculture, Missouri State University, USA;
[100], SeaWorld San Diego, [500], San Diego, California, USA;

[101], Department of Forensic Science, Environmental & Life Sciences, Trent University, Peterborough Ontario, Canada K[9]J [7]B[8];
[102], Shobhaben Pratapbhai Patel School of Pharmacy and Technology Management, SVKM'S NMIMS University, Mumbai, India;
[103], Dept. of Animal Science, University of Nebraska, Lincoln, USA;
[104], Species Preservation Laboratory, SeaWorld San Diego, California, USA;
[105], Biology Department, Acadia University, Wolfville, Nova Scotia, Canada B[4]P [2]R[6];
[106], Dept. of Pathobiology and Population Sciences, Royal Veterinary College, Hatfield, UK;
[107], School of Biology and Environmental Science, University College Dublin, Belfield, Dublin, Ireland;
[108], Animal Care and Science Division, John G. Shedd Aquarium, Chicago, Illinois, USA;
[109], School of Biology, The University of St Andrews, Fife, UK;
[110], Blizard Institute, Barts and the London School of Medicine and Dentistry, Queen Mary University of London, London, UK;
[111], Dept. of Pathology and Laboratory Medicine, David Geffen School of Medicine at UCLA, Los Angeles, USA;
[112], Center for Neurobehavioral Genetics, Jane and Terry Semel Institute for Neuroscience and Human Behavior, University of California Los Angeles, Los Angeles, CA, USA;
[113], Dept. of Genetics, Genomics and Informatics, University of Tennessee Health Science Center, College of Medicine, Memphis, TN, USA;
[114], Center for Computational and Genomic Medicine, Children's Hospital of Philadelphia, Philadelphia, USA;
[115], Rocky Mountain Biological Laboratory, Crested Butte, CO, USA;
[116], Dept. of Biochemistry, University of Otago, Dunedin, Otago, New Zealand;
[117], Department of Evolution, Ecology and Organismal Biology, The Ohio State University, Columbus, OH, United States; Translational Data Analytics Institute, The Ohio State University, Columbus, OH, United States;
[118], David Geffen School of Medicine, University of California, Los Angeles, CA, USA;
[119], Bioinformatics Interdepartmental Program, University of California, Los Angeles, CA, USA;Dept. of Biological Chemistry, University of California, Los Angeles, Los Angeles, California, USA;
[120], Fisheries and Oceans Canada, University Crescent, Winnipeg, Canada;Dept. of Biological Sciences, University of Manitoba, Winnipeg, Canada;
[121], School of Biological, Earth and Environmental Sciences, University of New South Wales, Sydney, Australia; Evolutionary Genetics Group, Department of Anthropology, University of Zurich, Zurich, Switzerland;
[122], Dept. of Human Genetics, David Geffen School of Medicine, University of California Los Angeles, Los Angeles, CA, USA;Dept. of Biostatistics, Fielding School of Public Health, University of California Los Angeles, Los Angeles, CA, USA;
[123], Division of Genetics & Metabolism, Oishei Children’s Hospital, Buffalo, New York, USA;
[124], Dept. of Drug Discovery and Biomedical Sciences, College of Pharmacy, University of South Carolina, SC, USA;Peromyscus Genetic Stock Center, University of South Carolina, SC, USA;
[125], Institute of Animal Reproduction and Food Research of Polish Academy of Sciences, Olsztyn, Poland;Institute for Veterinary Medicine, Nicolaus Copernicus University, Torun, Poland;
[126], Center for Tropical Research, Institute for the Environment and Sustainability, UCLA, Los Angeles, CA, USA;
[127], Dept. of Animal Science, College of Agriculture and Natural Resources, Laramie, Wyoming, USA ;
[128], Dept. of Preventive Medicine, University of Tennessee Health Science Center, College of Medicine, Memphis, TN, USA;Dept. of Genetics, Genomics and Informatics, University of Tennessee Health Science Center, College of Medicine, Memphis, TN, USA;

[129], Interdisciplinary Program in Genetics, Texas A&M University, College Station, Texas [77843], U
SA;
[130], Institute of Molecular Biology gGmbH, Mainz, Germany;Division of Molecular Embryology, D
KFZ-ZMBH Alliance, Heidelberg, Germany;
[131], Deutsches Krebsforschungszentrum, Division of Regulatory Genomics and Cancer Evolutio
n - B[270], Im Neuenheimer Feld [280], Heidelberg, Germany;
[132], Evolutionary Genetics Group, Department of Anthropology, University of Zurich, Zurich, Swit
zerland; Cetacean Ecology Research Group, School of Natural Sciences, Massey University, A
uckland, New Zealand; Global Ecology Partuyarta Ngadluku Wardli Kuu, College of Science an
d Engineering, Flinders University, Adelaide, Australia;
[133], Tikki Hywood Foundation, [7] Courtenay Road, Ballantyne Park, Harare, Zimbabwe;
[134], Center for Conservation Genomics, Smithsonian Conservation Biology Institute, Washington
, D.C., USA;
[135], Dept. of Biochemistry and Molecular Genetics, University of Illinois at Chicago, Chicago, IL,
USA;
[136], Dept. of Animal Sciences, University of Illinois at Urbana-Champaign, USA;
[137], Howard Hughes Medical Institute, Department of Neuroscience, University of Texas Southw
estern Medical Center, Dallas, Texas, USA;
[138], Neurogenetics of Vocal Communication Group, Max Planck Institute for Psycholinguistics, N
ijmegen, The Netherlands;
[139], Mother Infant Research Institute, Tufts Medical Center, Boston, MA, USA;Division of Obstetr
ics and Gynecology, Tufts University School of Medicine, Boston, MA, USA;
[140], Center for Neurobehavioral Genetics, Jane and Terry Semel Institute for Neuroscience and
Human Behavior, University of California Los Angeles, Los Angeles, CA, USA;Dept. of Psychiat
ry and Biobehavioral Sciences, David Geffen School of Medicine at UCLA, Los Angeles, CA, U
SA;
[141], Eli and Edythe Broad Center of Regenerative Medicine and Stem Cell Research, University
of California, Los Angeles, USA;
[142], Dept. of Pathology and Laboratory Medicine, University of Pennsylvania, Philadelphia, USA;
