## Supplementary material for "Divergent age-related methylation patterns in long and short-lived mammals": A workflow of using LUC version 1 clocks in mice

### A workflow to use Lifespan Uber Correlation (LUC version1) Clocks

Amin Haghani

01/15/2022

```
library(dplyr)
library(tidyr)
```

```
# import normalize beta values
miceadds::load.Rdata("normalized.Rdata", objname = "dat")



# import sample sheet
samples <- read.csv("samples.csv")





samples
```

```
## # A tibble: 6 x 3
##   Basename                Age Treatment
##   <chr>                 <dbl> <chr>
## 1 205600840027_R03C02 0.458 Drug
## 2 205619060031_R03C01 0.396 Drug
## 3 205619060041_R03C01 0.667 Vehicle
## 4 205619060072_R02C01 0.667 Vehicle
## 5 205619060077_R04C01 0.458 Drug
## 6 205619060077_R05C02 0.583 Vehicle
```

```
# import the LUC clock coefficients, as an example, I will only use the LUC panTissue clock
LUC_clock <- read.csv("LUC panTissue Coefficients.csv")

LUC_clock[1:8,]
```

```
##          CGid       Coef
## 1  Intercept -1.3737439
## 2 cg00059486 -0.4674907
## 3 cg00077617  0.1163220
## 4 cg00091964 -1.4065777
## 5 cg00272971  0.4034882
## 6 cg00281640 -0.3023834
## 7 cg00314427 -0.7086294
## 8 cg00374382 -0.5467259
```

```
## Prepare the data
dat <- dat %>% tibble::column_to_rownames("CGid") %>%
      dplyr::select(samples$Basename)%>%
```

```r
      # transpose the data
      t(.) %>% as.data.frame(.)%>%
      # add an intercept column
      mutate('Intercept'=1)%>%
      # select the LUC covariates
      dplyr::select(LUC_clock$CGid)

## Predict age and calculate age acceleration
samples <- samples %>%
  # predicting age
  mutate(epiAge = as.numeric(as.matrix(dat)%*%LUC_clock$Coef))%>%
  # calculate age acceleration
  mutate(AgeAccelation = as.vector(residuals(lm(epiAge~Age))))

samples[1:3,]
```

```
## # A tibble: 3 x 5
##   Basename               Age Treatment epiAge AgeAccelation
##   <chr>                <dbl> <chr>      <dbl>         <dbl>
## 1 205600840027_R03C02 0.458 Drug       0.261        -0.181
## 2 205619060031_R03C01 0.396 Drug       0.216        -0.101
## 3 205619060041_R03C01 0.667 Vehicle    0.532        -0.326
```

```r
## Analyzing the treatment effect using multivaraite linear regression
samples <- samples%>%
  # Convert the treatment variable to a dummy variable
  mutate(Treatment = as.numeric(as.character(
    factor(Treatment, levels = c("Vehicle", "Drug"),
           labels = c(0,1)))))

# The model design depends on the study design
lm.model <- lm(AgeAccelation~Treatment+Age, data = samples)
summary(lm.model)
```

```
##
## Call:
## lm(formula = AgeAccelation ~ Treatment + Age, data = samples)
##
## Residuals:
##        1        2        3        4        5        6
## -0.07702 -0.19539 -0.27607  0.12953  0.27241  0.14654
##
## Coefficients:
##             Estimate Std. Error t value Pr(>|t|)
## (Intercept)   2.0684     2.0986   0.986    0.397
## Treatment    -0.7161     0.6977  -1.026    0.380
## Age          -3.1779     3.2752  -0.970    0.403
##
## Residual standard error: 0.2786 on 3 degrees of freedom
## Multiple R-squared:  0.2599, Adjusted R-squared:  -0.2335
## F-statistic: 0.5267 on 2 and 3 DF,  p-value: 0.6367
```
